# Robust analysis of comparative subcellular omics with complex designs

**DOI:** 10.64898/2026.07.07.736960

**Authors:** Oliver M. Crook, Anna Andrejeva, Rayner M.L. Queiroz, Tom S. Smith, Lisa M. Breckels, Kathryn S. Lilley

## Abstract

Subcellular omics technologies now allow us to obtain insights into steady-state localisation and re-localisation of biomolecules in high-throughput. However, robust analysis of these experiments can be slow and challenging. Here, we show that existing approaches to differential localisation fall into two classes, marker-dependent and marker-free, that fail for fundamentally different statistical reasons, with failure modes that are not completely overlapping. We exploit this observation in SANDLE (statistical analysis of differential localisation experiments), which combines a marker-dependent generative model of subcellular niches with a marker-free regression test for changes in fractionation profiles. This dual strategy provides better control of false positives, up to 100-fold reduction in analysis time, and accommodates more complex experimental designs than existing methods. We demonstrate SANDLE’s versatility across a wide range of subcellular omics experiments, including drug-treatment responses, cross-species and life-cycle comparisons, post-translationally modified proteoforms, and RNA re-localisation. Co-analysing transcriptome and proteome dynamics during the unfolded protein response, we further reveal lncRNAs whose steady-state localisation is condition-dependent. By addressing key methodological limitations, SANDLE enables broad applications of subcellular omics from fundamental biology to clinical research.

## Introduction

Cellular compartmentalisation is fundamental to molecular function, with distinct organelles and subcellular niches providing specialised micro-environments for specific interactions (Gibson, 2009). The spatial organisation of biomolecules, including proteins and RNA, regulates access to interaction partners and cellular machinery, and many biological processes are controlled through dynamic relocalisation. Determining subcellular localisation is therefore a key step in understanding molecular function. Moreover, differential localisation in disease states may provide mechanistic insight into pathology by revealing disrupted spatial organisation (Kau *et al*., 2004; Davies, 2018).

To study steady-state localisation and dynamic re-localisation of biomolecules in high-throughput, one can couple gentle cell lysis and whole-cell fractionation with proteomics or transcriptomics, enabling simultaneous measurement of protein and RNA distributions across subcellular fractions (Christopher *et al*., 2021; Breckels *et al*., 2024; Villanueva *et al*., 2024). Further enrichment strategies can be included to study the localisation of post translationally modified proteoforms such as phosphorylation (Liu *et al*., 2021; Kafkia *et al*., 2022). A number of high-resolution subcellular maps have been generated using these technologies (Itzhak *et al*., 2016; Itzhak, 2017; Orre, 2019; Geladaki, 2019; Shin, 2020; Barylyuk *et al*., 2020; Mulvey, 2021; Moloney *et al*., 2023; Guérin *et al*., 2023; Christopher *et al*., 2025). Recent applications of these technologies have revealed condition-dependent localisation patterns, revealing how cellular perturbations influence subcellular protein and RNA dynamics (Villanueva *et al*., 2024). This approach has also enabled comparative analyses across different biological contexts, as demonstrated by studies examining protein localisation changes across life-cycle stages and species in African Trypanosomes (Moloney *et al*., 2023).

While computational methods for analysing protein localisation dynamics are well established, equivalent statistical frameworks for analysing RNA relocalisation from fractionation-based experiments remain limited, motivating unified approaches that can analyse both molecular modalities. Fractionation-based spatial transcriptomics approaches have also been developed to characterise RNA localisation across subcellular compartments, including CeFra-seq and related methodologies (Benoit Bouvrette *et al*., 2018; Adekunle and Wang, 2020). These studies demonstrated that transcripts exhibit reproducible and asymmetric distribution patterns across biochemical fractions, analogous to spatial proteomics datasets. However, dedicated statistical frameworks for detecting differential RNA localisation across conditions remain limited, and most analyses rely on clustering, correlation-based comparisons, or descriptive profiling. This highlights the need for unified methods capable of quantifying RNA relocalisation with statistical control, alongside protein and post-translational modification datasets.

Analysis of subcellular-omics data typically employs supervised machine learning approaches, where marker proteins with well characterised localisations guide the classification of proteins to specific subcellular niches (Crook *et al*., 2018, 2020). Several computational methods have been developed to assess protein dynamics including the Movement-Reproducibility (MR) method (Itzhak *et al*., 2016), mobility score (MS) (Martinez-Val *et al*., 2021), TRANSPIRE (Kennedy *et al*., 2020), BANDLE (Crook *et al*., 2022) and TransGCN (Wang *et al*., 2024). While MR and MS methods do not rely on marker proteins, TransGCN and TRANSPIRE leverage marker profiles to generate synthetic training data for graph convolutional networks and Gaussian processes, respectively. BANDLE incorporates marker profiles through a hierarchical Bayesian mixture model. Independent benchmarking has demonstrated that TransGCN, followed by BANDLE, achieves the highest accuracy in determining differential localisations (Wang *et al*., 2024).

Current methodological approaches, while powerful, face several key limitations when analysing complex experimental designs. These include challenges in comparing proteins across species where direct protein mapping may not be possible, and dependence on extensively curated marker proteins which may be limited in some systems (Christopher *et al*., 2021). Additionally, existing methods struggle to accommodate experimental designs with varying numbers of conditions (such as drug concentration series) or uneven numbers of replicates between control and disease states. As proteomics coverage and experimental complexity continue to expand, there is a growing need for approaches that are both computationally efficient and scalable while maintaining strict control over false positives to prevent resource-intensive follow-up studies.

Here we present SANDLE (Statistical Analysis of Differential Localisation Experiments), which implements this dual-threshold strategy in a fast, modular framework. We show that SANDLE achieves near-zero false positive rates at standard thresholds where TransGCN and BANDLE produce 50% and 10% false positives respectively, completes analyses in seconds rather than the minutes to hours required by current methods, and accommodates experimental designs that existing tools cannot handle for example paired designs. We further apply SANDLE to PhosphoLOPIT, a new spatial phosphoproteomics dataset we generated in U-2 OS cells, identifying interface disruption and phosphatase recognition as the dominant features distinguishing phosphorylation-dependent localisation events. Application to African Trypanosomes reveals life-cycle stage and species specific localisation, included differential localisation of orthologous glycolytic enzymes. Further, we applied SANDLE to a multi-omic (transcriptome and proteome) subcellular dataset revealing complex changes in the proteome and transcriptome, including re-localisation of lncRNAs, during the unfolded protein resposne. SANDLE is made available in the BANDLE Bioconductor package.

## Results

### Development of the SANDLE method

During SANDLE’s development we identified that marker-dependent and marker-free approaches to differential localisation analysis fail for fundamentally different statistical reasons. Marker-dependent methods generate false positives when marker distributions are noisy or poorly reproducible across replicates, since the inferred niche distributions are themselves uncertain. Marker-free methods generate false positives when the biochemical properties of subcellular niches shift between conditions (Crook *et al*., 2022), altering fractionation patterns globally without any genuine re-localisation. Crucially, these two error sources are largely independent: a noisy marker problem does not produce a coherent global fractionation shift, and a global fractionation shift does not selectively perturb marker variance. This observation motivates a simple but unexplored strategy: exploit evidence from both marker-dependent and marker-free approaches to determine a differential localisation event. The conjunction rule eliminates both classes of false positive, since genuine re-localisation events typically generate evidence under both criteria.

SANDLE’s main innovation is to exploit a combination of marker-dependent and marker-free approaches to achieve robust differential localisation analysis. In the marker-dependent component, we fit multivariate t-distributions to marker protein profiles for each subcellular niche across replicates and conditions. These fitted distributions enable calculation of differential localisation probabilities by quantifying how well a protein’s profile matches different niches under varying conditions. Complementarily, the marker-free approach employs regression analysis on protein profiles, testing for significant interaction effects between subcellular fractions and conditions. This generates both a statistical p-value and a difference in residual sum of squares, providing an orthogonal metric for ranking localisation changes. Differential localisation probability reflects the confidence that a protein changes compartmental identity, whereas the FDR reflects statistical evidence for profile differences; these capture distinct aspects of spatial reorganisation.

By implementing dual thresholds across both approaches, SANDLE effectively filters out false positives that would otherwise arise from the limitations of each method in isolation. This hybrid strategy addresses two key sources of error: noisy marker distributions in marker-dependent methods and biochemical property changes in marker-free approaches. The resulting methodology provides both statistical confidence through p-values and an intuitive effect size metric for ranking localisation changes. The insight that marker-dependent and marker-free approaches fail in fundamentally different ways led to SANDLE’s novel dual-threshold strategy, effectively eliminating both sources of false positives. Rather than choosing between marker-based classification and profile regression, SANDLE leverages the complementary strengths of both approaches to achieve superior accuracy.

To further increase the utility of SANDLE, we expanded the range of applicable subcellular experiments by allowing incorporation of paired design, varying numbers of conditions and varying number of replicates, which are frequent during clinical studies. This allows a wide set of experiments to be performed using SANDLE.

### SANDLE is fast and robust

To evaluate SANDLE’s effectiveness, we tested its ability to detect differential localisation across 60 simulated experiments with known ground truth under diverse scenarios (see methods). We assessed performance using the area under the receiver operating characteristic curve (AUC), comparing false positive rates (FPR) against true positive rates (TPR). While the marker-free and marker-dependent modules individually achieved AUCs of approximately 92.5, their combination improved performance to an AUC of 95, demonstrating the synergistic benefit of this hybrid approach (Fig. 1A).This improvement arises because the two components capture orthogonal failure modes, reducing both marker-driven noise and profile-driven artefacts. Comparative analyses revealed SANDLE’s superior performance against both TransGCN (Wang *et al*., 2024) (Fig. 1B) and BANDLE (Crook *et al*., 2022) (Fig. 1D), with BANDLE slightly outperforming TransGCN (Fig. 1C).

**Figure 1:**
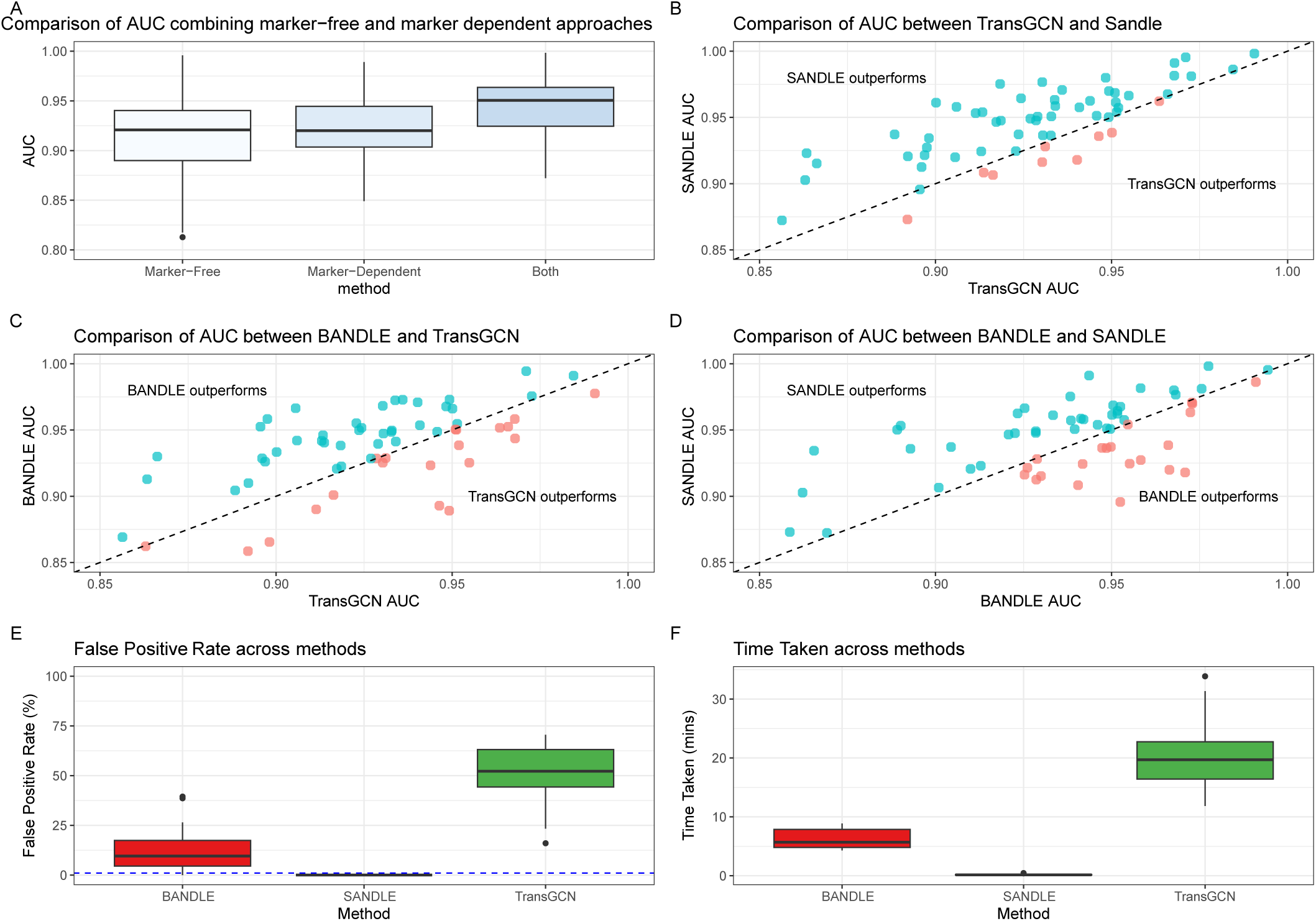
Comparison of differential localisation methods. A) Area under curve (AUC) of receiver operating characteristics (ROC) curves across simulations. 60 simulations are considered across a range of scenarios (see methods). B) SANDLE AUCs against TransGCN AUCs, each pointer represents evaluation on the same dataset. C) As for B but with BANDLE against TransGCN. D) As for B but for SANDLE against BANDLE E) Boxplots of distribution of false positive rates at nominal thresholds for each method across simulation scenarios. Dotted line indicated 1% false discovery rate. F) Clock times for each approach with distribution across simulation scenarios. All boxplots are in the style of Tukey.

We further evaluated method performance using each algorithm’s recommended thresholds for reliable prediction: FDR of 0.01 for TransGCN, differential localisation probability of 0.99 for BANDLE, and combined thresholds of 0.99 probability and 0.01 FDR for SANDLE. At these thresholds, SANDLE achieved near-zero false positives, while BANDLE and TransGCN showed approximately 10% and 50% false positive rates, respectively. TransGCN’s high false positive rate suggests poorly calibrated FDR estimation, which could lead to wasted resources in validation experiments. Notably, SANDLE completed analyses in under 11 seconds on average, substantially faster than both BANDLE (>5 minutes) and TransGCN (20 minutes) (Fig. 1F), demonstrating that SANDLE’s improved robustness does not come at the cost of computational efficiency. The 100-fold reduction in analysis time enables complex experimental designs that were previously intractable such as clinical cohorts and large ligand screens.

While AUC provides a broad performance metric, practical applications require selecting specific thresholds to prioritise candidates for experimental validation. We therefore evaluated method performance using standard thresholds across multiple simulated scenarios and a relaxed threshold for SANDLE. TransGCN, while highly sensitive, frequently generated more false positives than true positives (Fig. S1 A), BANDLE demonstrated similar sensitivity with substantially reduced false positives (Fig. S1 B). At standard thresholds, SANDLE effectively eliminated false positives, though at some cost to sensitivity (Fig. S1 C). However, using slightly relaxed thresholds (probability: 0.95 and FDR: 0.05), SANDLE maintained minimal false positives while achieving improved sensitivity (Fig. S1 D), offering a practical balance for experimental validation.

Beyond improved statistical performance, SANDLE substantially expands the range of experimental designs that can be analysed within a single statistical framework (Table 1). In addition to conventional two-condition comparisons, SANDLE accommodates paired designs, unequal replicate numbers, multiple experimental conditions, cross-species comparisons, and peptide- and transcript-level localisation analyses without requiring separate analysis pipelines. These practical capabilities

**Table 1:** Comparison of practical capabilities of BANDLE and SANDLE from the perspective of an experimental user. *Possible by running BANDLE between pairs but doesnt exploit shared variance structure. † would require running BANDLE on subsets of pairs.

| Capability | BANDLE | SANDLE |
| --- | --- | --- |
| Two-condition comparison | ✓ | ✓ |
| Unequal replicate numbers | — | ✓ |
| Paired experimental designs | * | ✓ |
| More than two conditions | † | ✓ |
| Cross-species comparison | — | ✓ |
| RNA localisation analysis | — | ✓ |
| PTM localisation analysis | — | ✓ |
| Protein-level localisation | ✓ | ✓ |
| Probabilistic localisation assignments | ✓ | ✓ |
| Regression-based statistical testing | — | ✓ |
| Runtime (typical dataset) | hours-days | seconds-minutes |

### The effect of experimental design on SANDLE

Having established SANDLE’s robust performance, we next investigated how experimental design parameters influence its sensitivity and specificity. Understanding these relationships is crucial for optimising experimental designs while managing resource constraints. We systematically evaluated the impact of fraction resolution and replicate number by analysing a simulated 10-fraction experiment, selectively averaging neighbouring fractions and varying replicate numbers between 2 and 5 (see methods).

As expected, increasing either the number of subcellular fractions or replicates improved SANDLE’s performance, enhancing true positive detection(Fig. 2A and 2D) while reducing false positives (Fig. 2B). To quantify these relationships, we used a linear model that revealed the relative contribution of each parameter. The combined marker-free and marker-dependent approach consistently improved AUC, with each additional replicate contributing an AUC increase of approximately 0.012 and each additional fraction contributing 0.007 (Fig. 2C). These quantitative relationships provide practical guidance for experimental design, allowing researchers to optimise the trade-off between experimental complexity and statistical power while considering available resources and technical constraints.

**Figure 2:**
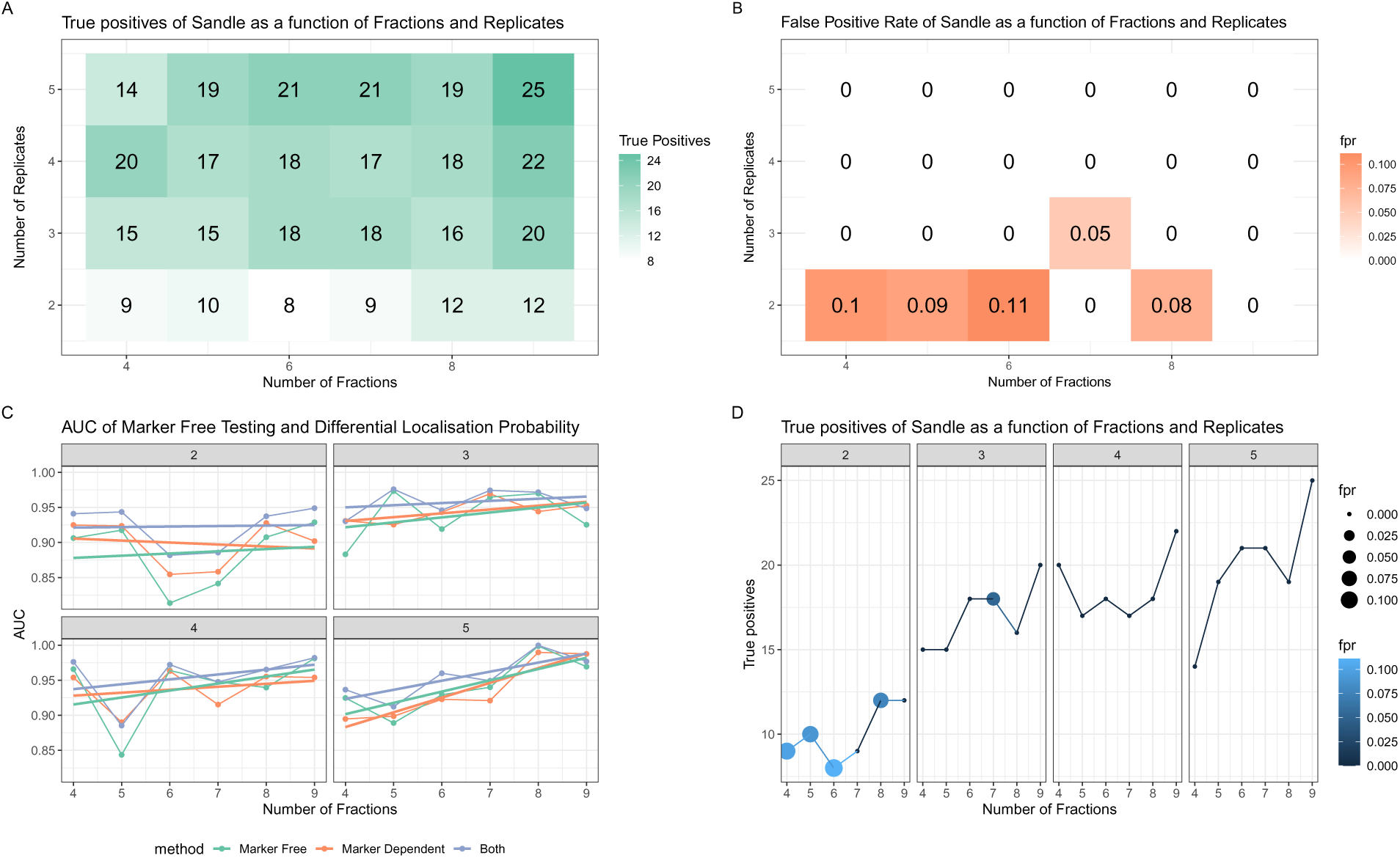
Experimental parameters influence on SANDLE. A) True positives (max 30) as a function of the number of subcellular fractions and number of replicates. B) False positive rate as a function of fractions and replicates. C) AUC as a function of fractions and replicates (facets) and colours indicate different methods. D) True positives as a function of fractions and replicates (facets) point-size and colour also indicates false positive rate (fpr).

### SANDLE recovers experimentally validated differential localisations

To validate SANDLE’s performance on real biological data, we analysed three previously published MS-based subcellular proteomics datasets spanning distinct biological contexts: cell-type comparison (antibody-producing CHO-K1 versus plasma cell-derived MPC-11 cells) (Kretz *et al*., 2022), insulin-induced changes in adipocytes (Conway *et al*., 2026), and drug response (HCC827 cells treated with the EGFR inhibitor gefitinib) (Orre, 2019). Each dataset presents distinct challenges and includes well-characterised protein re-localisation events.

Analysis of the cell-type comparison experiment using SANDLE (FDR ≤ 0.05, differential localisation probability ≥ 0.95) identified 194 proteins with altered subcellular distributions between CHO-K1 and MPC-11 cells. Gene Ontology enrichment analysis revealed significant enrichment of protein trafficking-related terms (Supplementary Fig. 2). Notably, multiple components of the exocyst complex (EXO3, EXO4, and EXO8), together with their regulator RAB8A, showed high-confidence differential localisation, suggesting that altered trafficking pathways may contribute to differences in antibody production capacity. SANDLE also recovered previously validated differential localisations, albeit with varying confidence: Eif2ak3 (FDR = 0.037, DL Prob = 0.36), Eif4g1 (FDR = 1.7 × 10^−5^, DL Prob = 0.80), Rab5b (FDR = 0.0087, DL Prob = 0.997), and Stx17 (FDR = 5.59 × 10^−9^, DL Prob = 0.914). The comparatively low confidence for Eif2ak3 is consistent with prior western blot analyses of subcellular fractions, which showed similar profiles between cell lines for both Eif2ak3 and the ER marker Calr (Kretz *et al*., 2022). This suggests that earlier classifications may have been influenced by outlying MS measurements (Fig. 3B).

**Figure 3:**
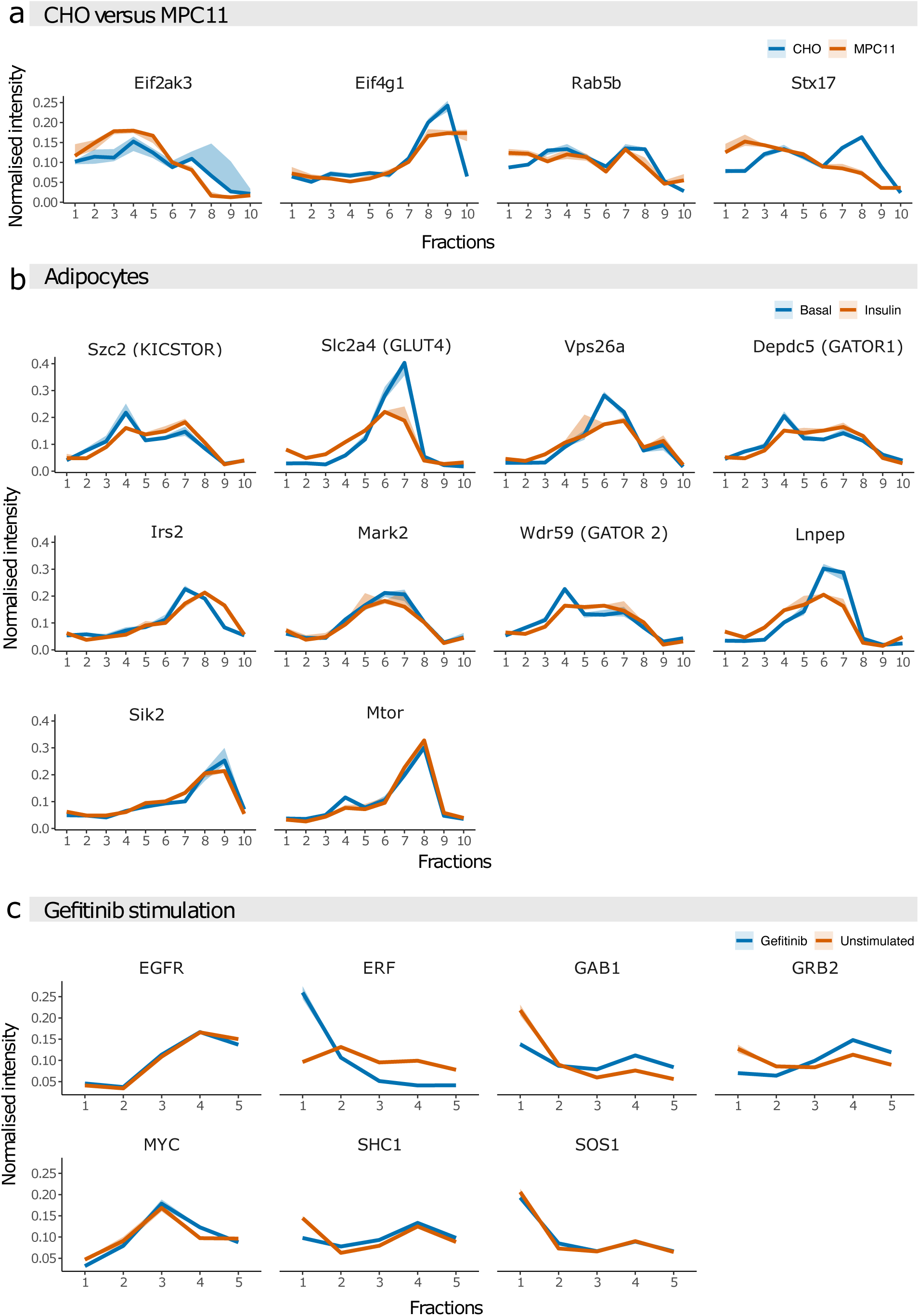
Analysis of well studied differential localisations. A) Example differential localisations in the cell-type comparison case study, note outlying measurement in top left panel. Protein intensity is plotted as a function of subcellular fraction. B) As for A but example differential localisations identified in adipocytes experiment C) As for B but example differential localisations identified in Gefitinib treatment. All ribbons represent standard errors from n = 3 replicates, except panel C which is 2 replicates

**Figure 4:**
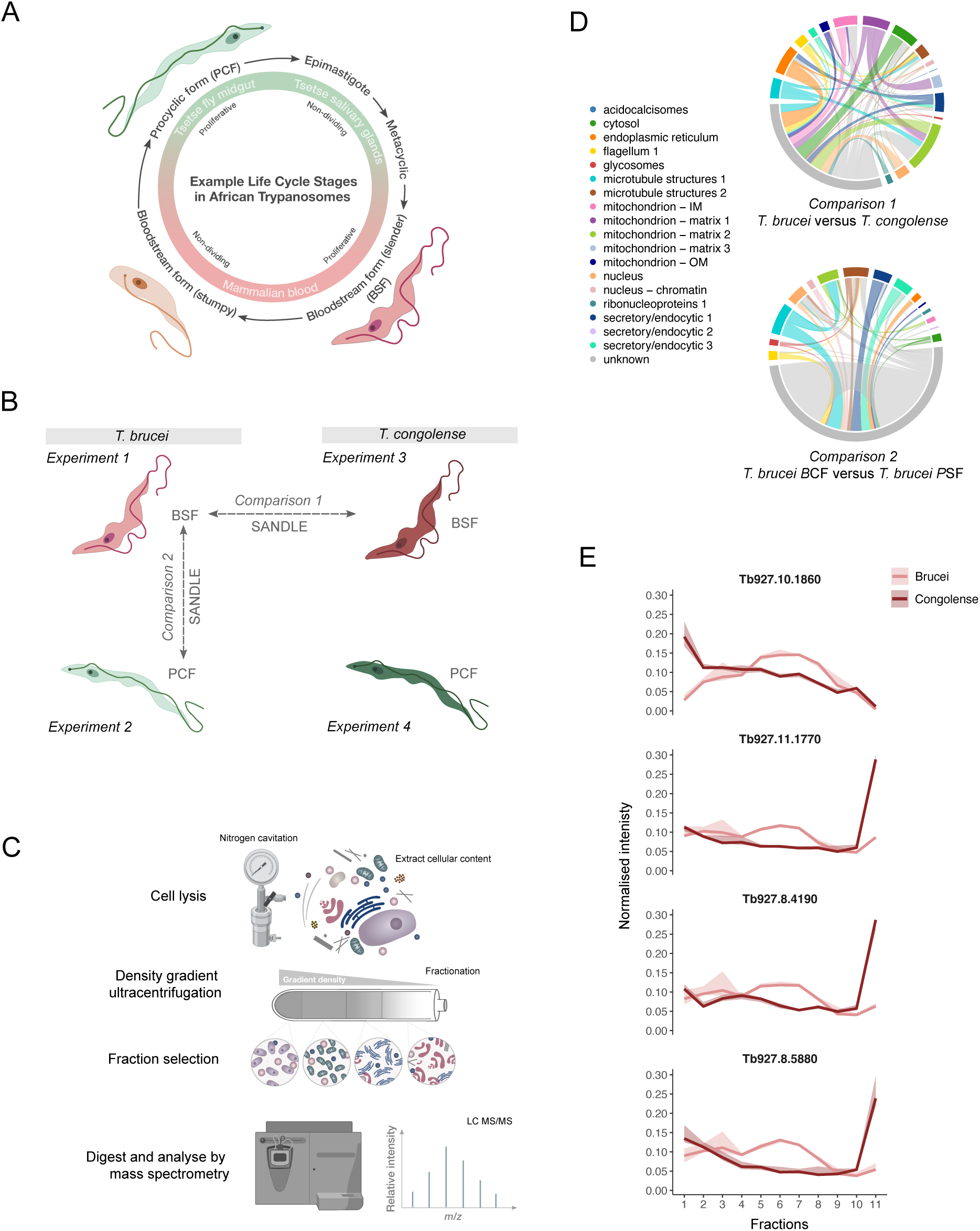
Comparative analysis between life stages and species. A) Example life cycle stages in African Trypanosomes B) diagram representing comparisons draw in this analysis C) Experimental protocol for HyperLOPIT experiments. D) Alluvial diagrams showing differential localisations between comparisons. E) Ribbon diagrams (standard errors) showing gradient profiles of referenced protein from n = 3 replicates. *T. brucei* protein Tb927.10.1860 is a glycosomal protein by microscopy therefore indicating a glycosomal profile.

Conway *et al*. (2026) performed subcellular proteomics on 3T3-L1 adipocytes, including an additional fraction enriched for lipid droplet-associated proteins. In this system, SANDLE identified previously implicated proteins such as GLUT4 (P14142; FDR = 1.5 × 10^−3^, DL Prob = 0.775; Fig. 3B). In addition, the uncharacterised protein C3ORF18 was identified with strong support (Q8BGK9, FDR = 2.0 × 10^−4^, DL Prob = 0.91), providing further evidence for its insulin-dependent role as discussed in Conway *et al*. (2026). At thresholds of differential localisation probability ≥ 0.8 and FDR *<* 0.05, SANDLE identified 69 proteins with altered localisation. KEGG pathway enrichment analysis confirmed over-representation of insulin signalling components (FDR = 0.0026), indicating that SANDLE captures both established and potentially novel regulators of insulin response.

We next analysed the EGFR inhibition experiment. EGFR, ERF, and SOS1 were excluded as markers used in the original study, as they were expected to undergo differential localisation (Hyatt and Ceresa, 2008; le Gallic *et al*., 1999; Saha *et al*., 2012). In total, 165 proteins showed evidence of differential localisation (FDR *<* 0.01, differential localisation probability ≥ 0.95; Supplementary Fig. 3). Several proteins with prior evidence of re-localisation were supported by SANDLE (Table 2). Interestingly, EGFR and MYC exhibit strong FDR support but comparatively low differential localisation probabilities, suggesting more complex or heterogeneous spatial reorganisation rather than simple compartmental shifts. In contrast, key EGFR signalling components GRB2, GAB1, and ERF show strong evidence across both metrics. Fig. 3D highlights the range of re-localisation magnitudes detected, from subtle (MYC) to pronounced (GRB2) spatial changes following EGFR inhibition.

**Table 2:** Differential localisation results showing false discovery rate (FDR) and differential localisation (DL) probability for selected proteins.

| Protein | FDR | DL Probability |
| --- | --- | --- |
| EGFR | $3.86 \times 10^{-5}$ | 0.1755 |
| ERF | $1.70 \times 10^{-6}$ | 0.9165 |
| GRB2 | $3.75 \times 10^{-5}$ | 0.9915 |
| GAB1 | $5.75 \times 10^{-5}$ | 0.8910 |
| MYC | $4.33 \times 10^{-2}$ | 0.2710 |
| SHC1 | $1.38 \times 10^{-4}$ | 0.7735 |
| SOS1 | $8.61 \times 10^{-2}$ | 0.6970 |

**Table 3:**
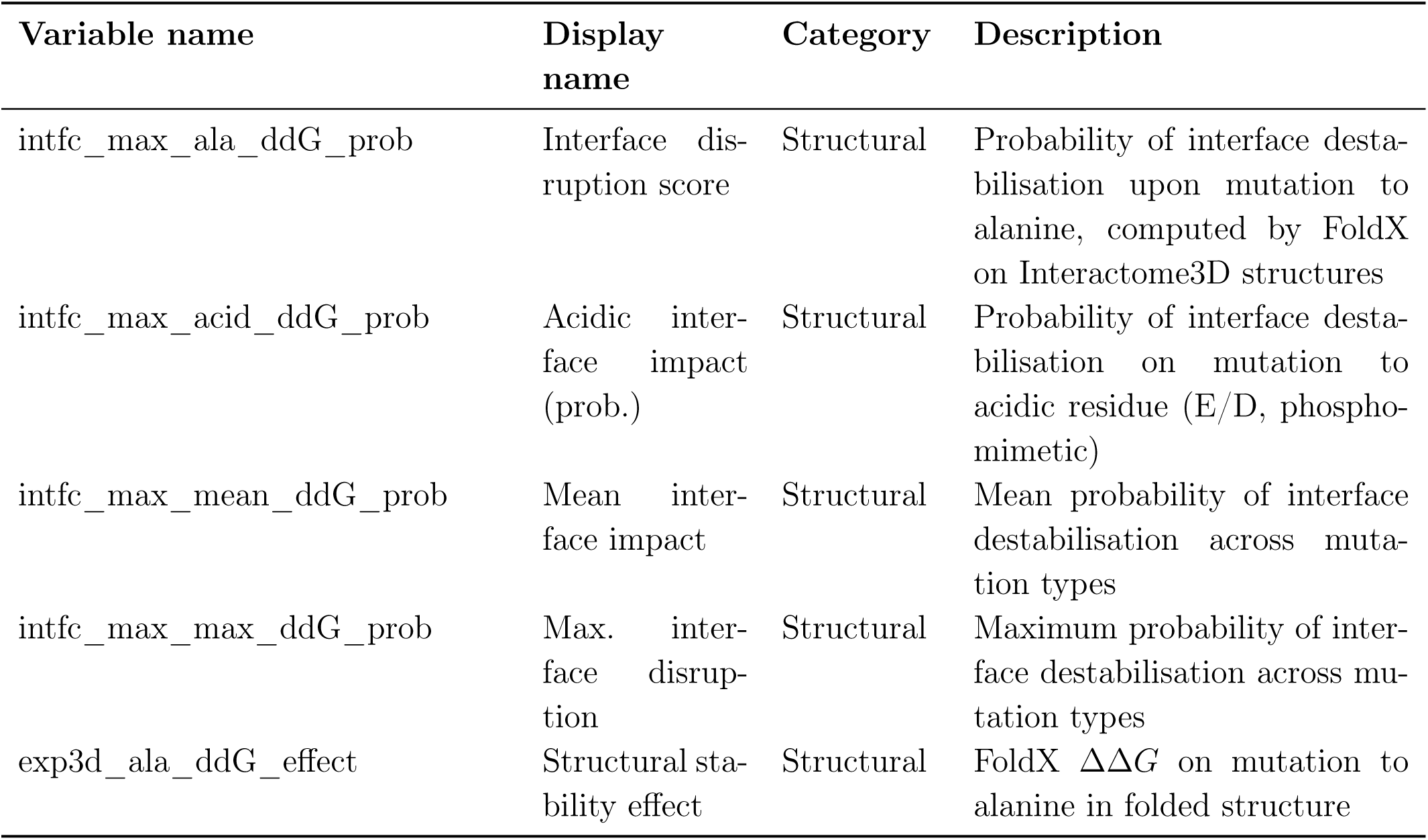
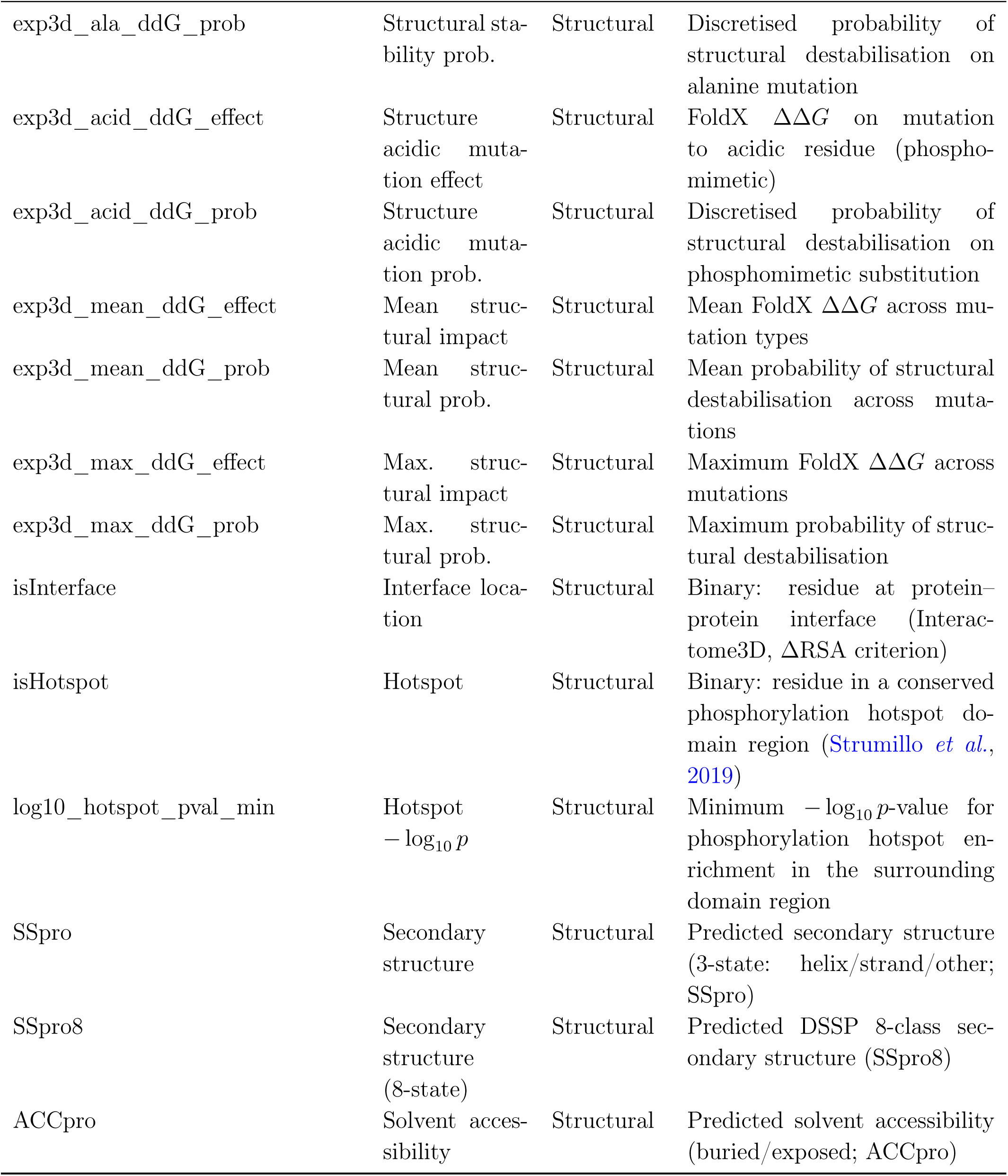
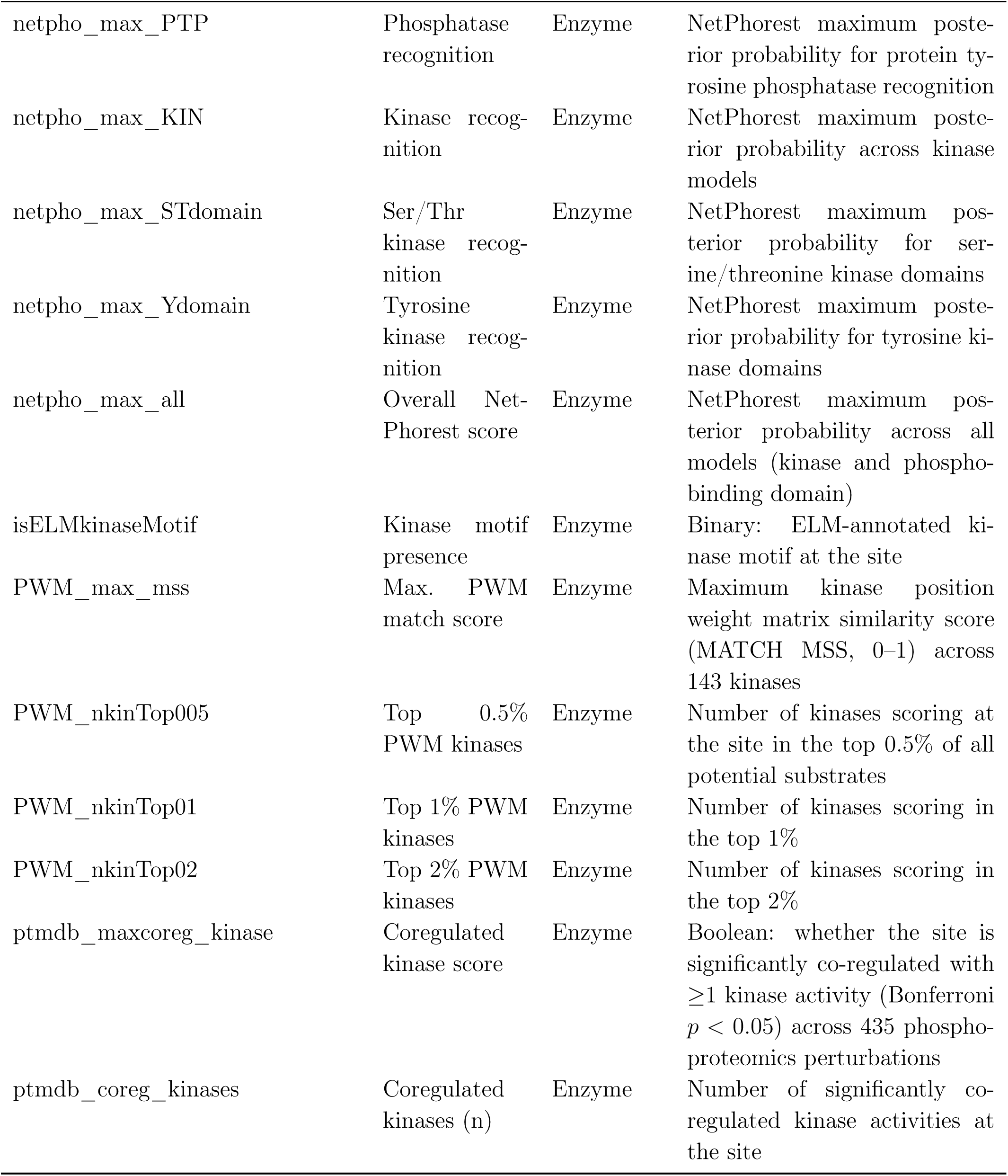
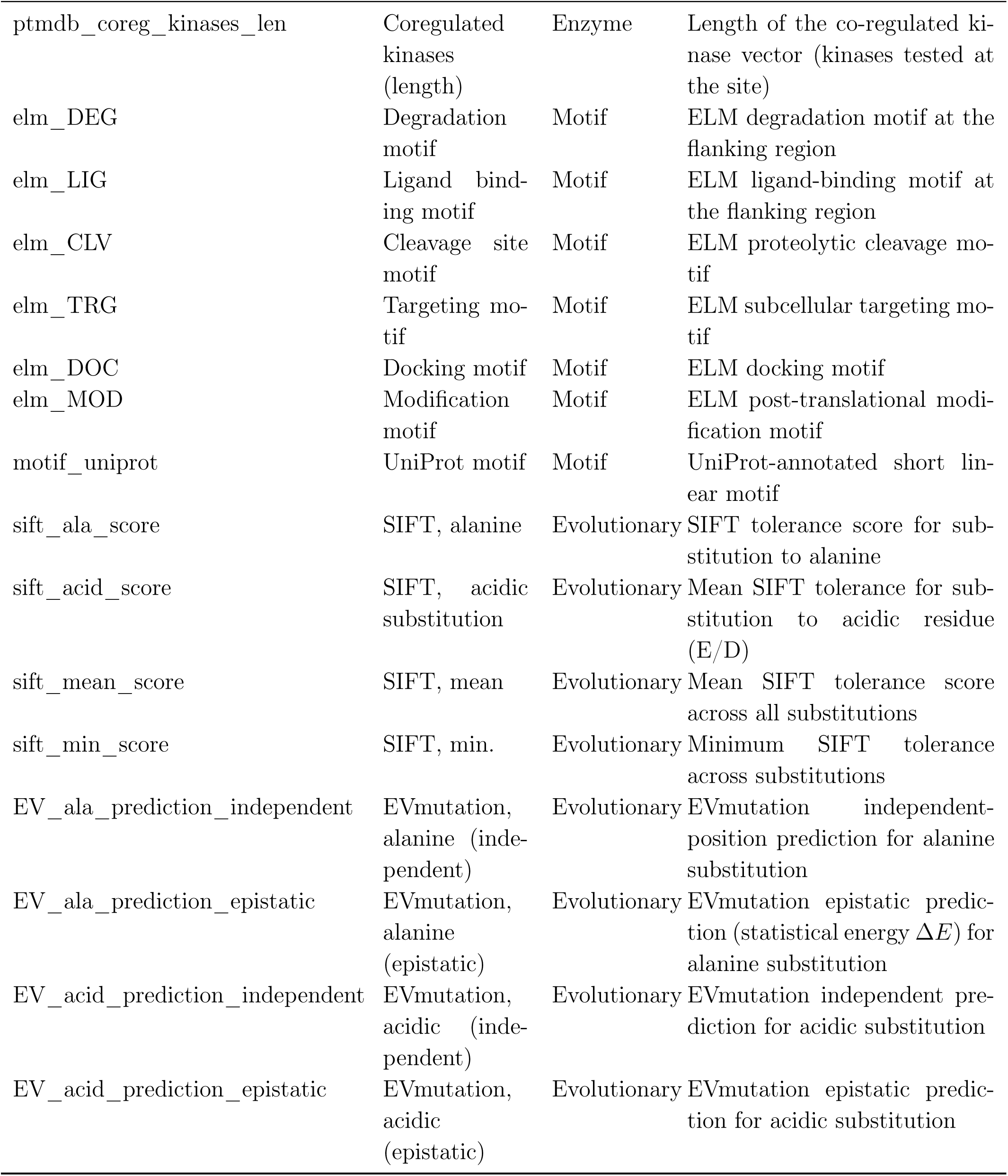
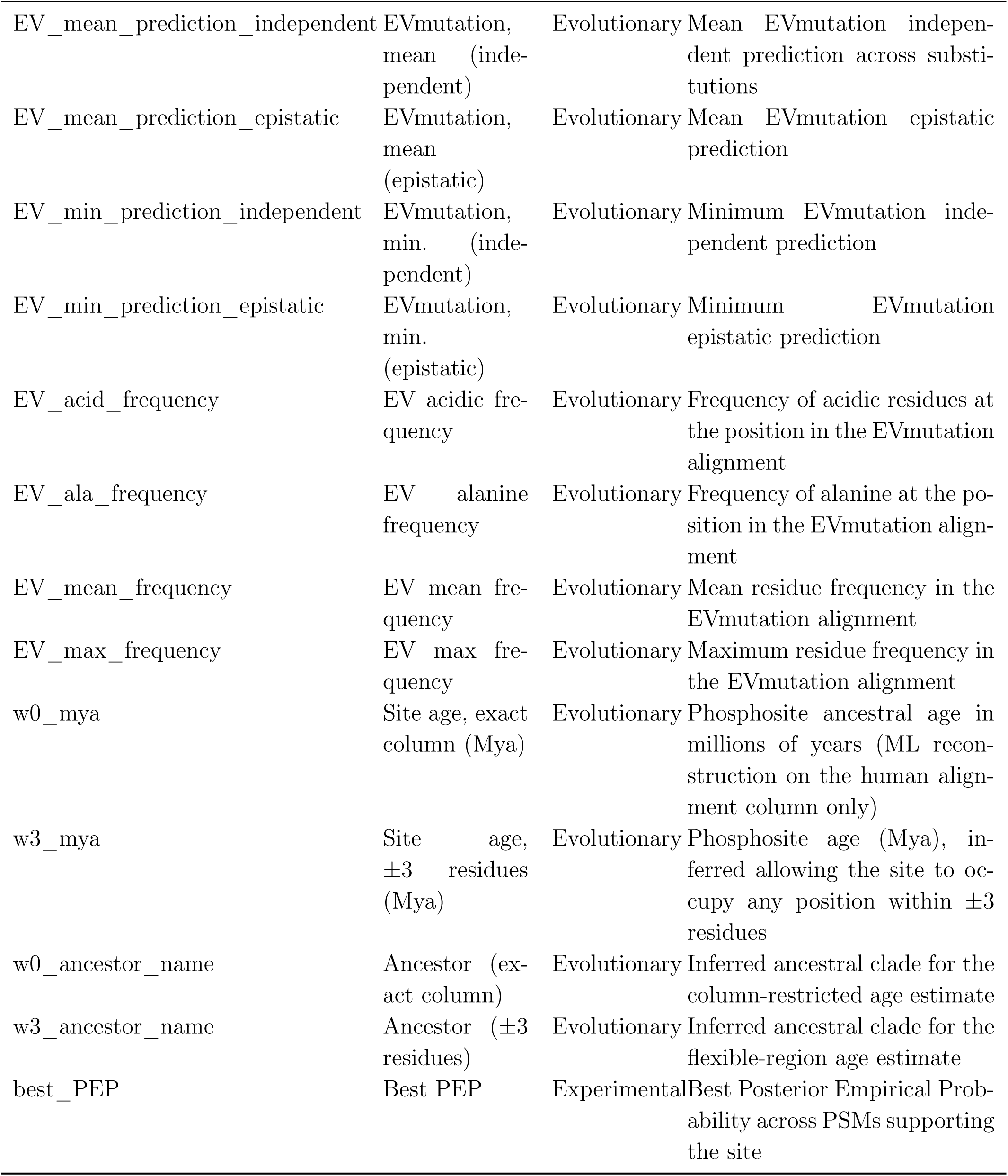
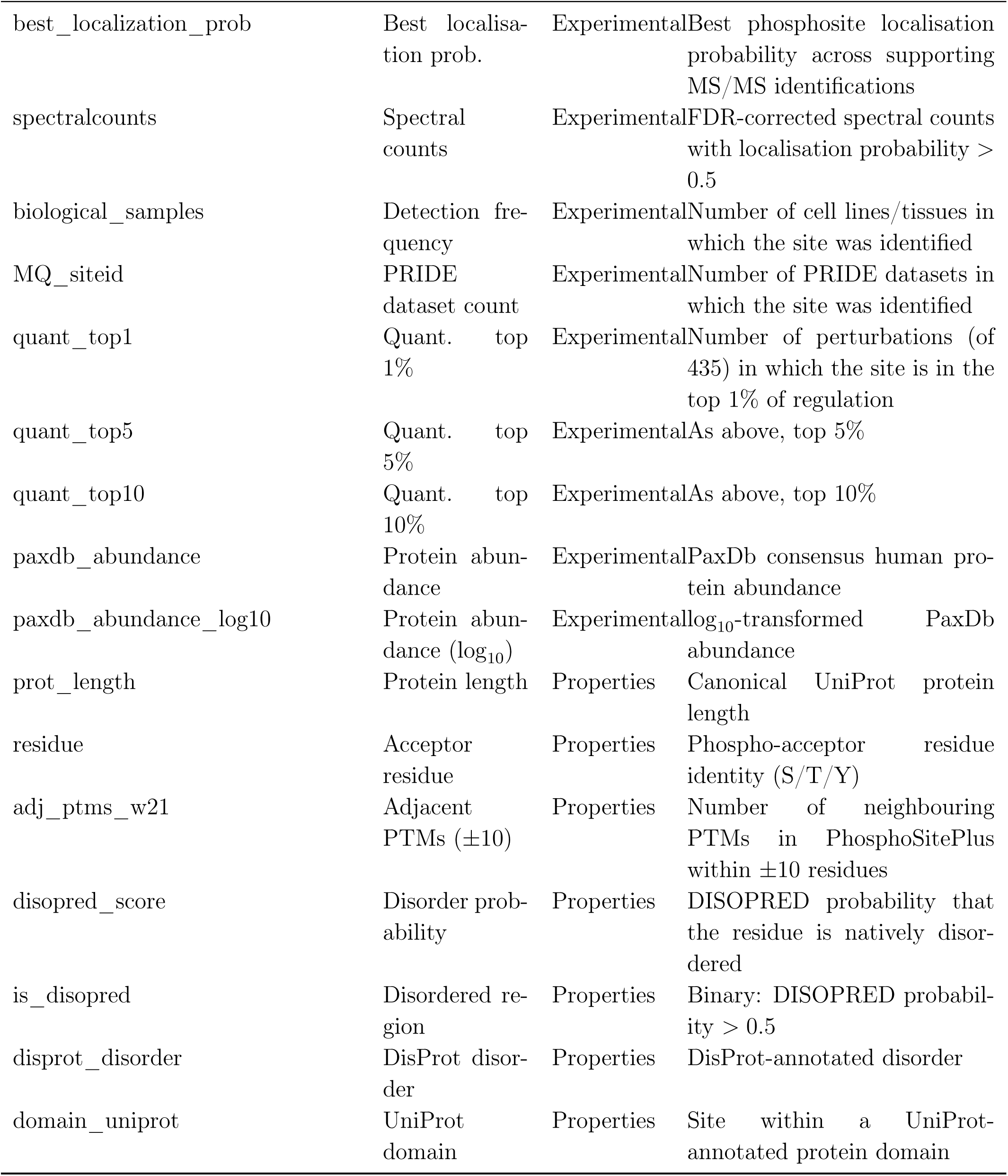
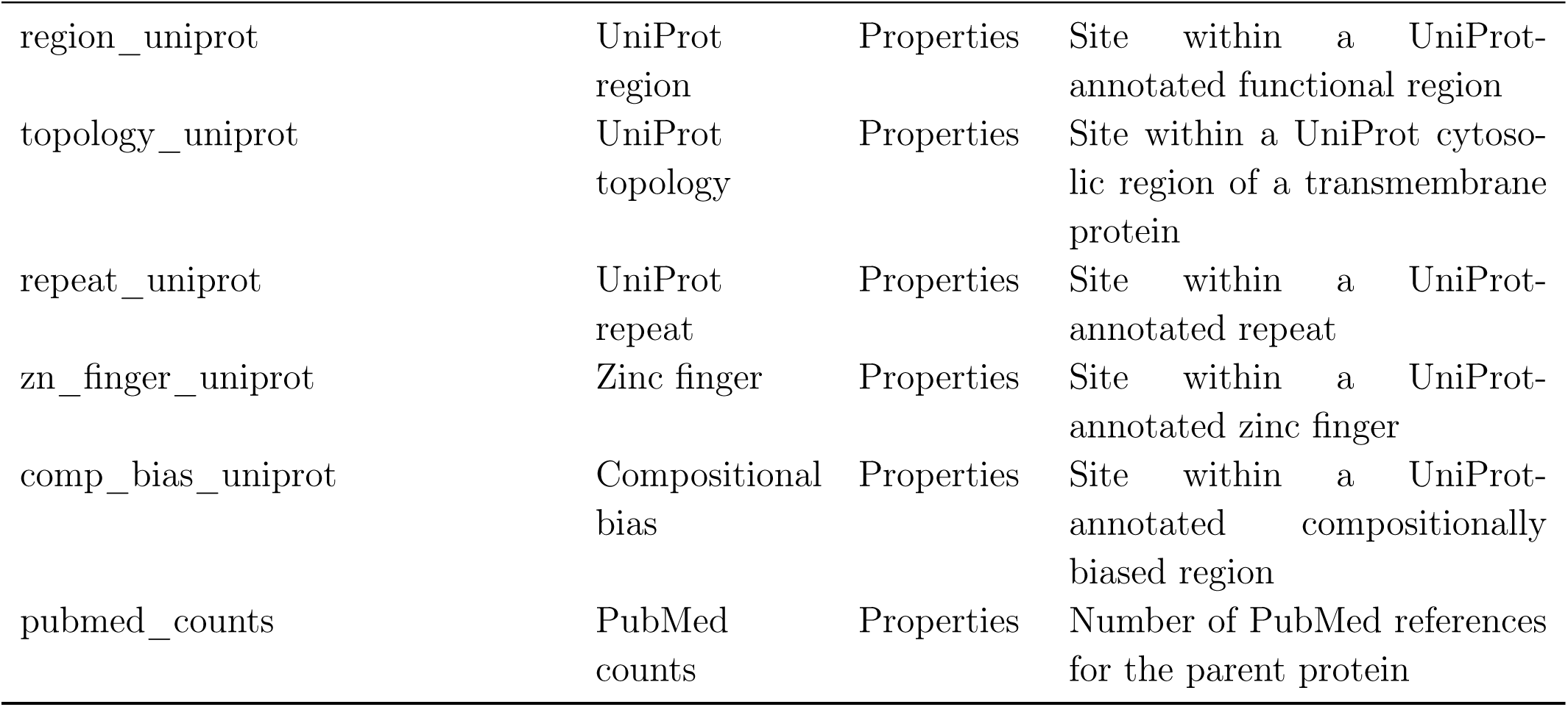
Phosphosite-level features used to evaluate associations with phosphorylation-dependent localisation, grouped by functional category. Variable names match the analysis code; display names match Figure 5E where applicable. Definitions are based on Ochoa *et al*. (2020) Supplementary Note 1.

### SANDLE enables cross-species and life-cycle analysis of protein localisation in Trypanosomes

To demonstrate SANDLE’s ability to handle complex experimental designs, we analysed protein localisation across species and life-cycle stages in Trypanosomes (Moloney *et al*., 2023). This analysis presents multiple technical challenges: mapping proteins between species (*T. brucei* and *T. congolense*) and comparing different life stages within *T. brucei*. We re-analyse previously published datasets that captured the spatial proteomes of bloodstream and procyclic forms in both species, providing an ideal test case for SANDLE’s comparative capabilities. While previous analysis (Moloney *et al*., 2023) relied on discrete classification differences to assess changes, potentially missing subtle re-localisation events, SANDLE’s statistical framework enables detection of more nuanced spatial proteome dynamics.

First, we applied SANDLE to compare life-cycle stages. SANDLE identified 76 proteins with differential localisation between life cycle stages, with the majority showing changes between poorly defined localisation patterns. However, several proteins displayed clear and biologically interesting localisation changes. For example, the uncharacterised protein Tb927.11.5200, which is conserved across kinetoplastids, shows distinct nuclear localisation in bloodstream form but redistributes to the cytosol in procyclic form. This protein contains a strong C-terminal basic motif (KRKRRGGRGE), which NLStradamus (Nguyen Ba *et al*., 2009) computationally predicts as a high-confidence nuclear localization signal, aligning with its observed nuclear localization in bloodstream form. Another notable example is phospho-glycerate kinase (PGK), where the data suggest a mixed and unclear localisation patterns in bloodstream form that re-distributes, according to subcellular proteomics, toward stronger cytosolic enrichment in procyclic form and an uncertain localisation in imaging of the tagged protein (Billington *et al*., 2023). In bloodstream forms, where glucose is the primary energy source, glycolysis is compartmentalised within glycosomes and PGK is therefore expected to localise predominantly to this organelle. In contrast, procyclic parasites rely primarily on L-proline metabolism and oxidative mitochondrial pathways, reducing their dependence on glycosomal glycolysis (Smith *et al*., 2017). The altered fractionation profile observed by SANDLE is therefore consistent with changes in the balance between glycosomal and cytosolic pools of PGK, reflecting broader metabolic remodelling during life-stages.

For cross-species comparison between *T. brucei* and *T. congolense*, analysis was restricted to one-to-one orthologs (Emms and Kelly, 2019) to ensure direct comparability, and we employed only SANDLE’s marker-free module as marker protein behaviour may not be conserved between species. Given the additional complexity of cross-species comparison, we implemented a stricter significance threshold (FDR < 0.0001). This approach identified 404 proteins with significant localisation differences between species, where fractionation profiles indicated distinct subcellular niches. The substantial number of differences, even under these stringent criteria, suggests considerable evolutionary divergence in protein localization between these closely related species.

Examining the data more closely revealed an intriguing set of proteins showing glycosomal localisation in *T. brucei* but different patterns in *T. congolense*. Glycosomes are peroxisome-related organelles characteristic of kinetoplastid parasites that compartmentalise glycolytic enzymes and other metabolic pathways essential for parasite energy metabolism. These observations are biologically plausible because *T. brucei*, particularly in its bloodstream stage, relies heavily on glycosomal glycolysis, whereas *T. congolense* exhibits greater dependence on mitochondrial metabolism and succinate production (Steketee *et al*., 2021). Fractionation profiles for these proteins are highly correlated in *T. brucei*, with one previously confirmed as glycosomal in microscopy (Tb927.10.1860) (Billington *et al*., 2023). While microscopy validation in *T. brucei* has been challenging due to technical limitations, the strong correlation of MS profiles supports a glycosomal localisation. The coordinated redistribution of *T. congolense* profiles suggests genuine species-specific differences in organelle targeting, potentially reflecting adaptations to different host niches and metabolic requirements.

### SANDLE enables subcellular analysis of protein post-translational modifications

A central question in cell biology is how post-translational modifications regulate subcellular localisation, and phosphorylation is the most prevalent and dynamically regulated modification in eukaryotes. To enable systematic analysis of phosphorylation-dependent localisation, we developed PhosphoLOPIT, a spatial phosphoproteomics workflow that pools subcellular fractions prior to phosphopeptide enrichment, ensuring that phosphorylated and unmodified peptide measurements derive from identical underlying fractions and avoiding the systematic enrichment biases that confound separate-fraction approaches. Previous studies have mapped protein succinylation, a modification that occurs primarily in mitochondria and links to metabolic regulation, across subcellular compartments (Kafkia *et al*., 2022). Here we applied PhosphoLOPIT to U2-OS cells but previous studies lacked a statistical framework for peptide-level analysis of subcellular proteomics data. SANDLE enables this analysis directly. Breifly, subcellular fractions were labelled with TMT reagents and pooled prior to enrichment for modified peptides using titanium dioxide for phosphorylated peptides, ensuring that phosphorylation measurements were derived from identical underlying subcellular fractions (Fig. 5 A). This design avoids enriching each fraction separately, which can introduce systematic biases due to differences in peptide complexity and enrichment efficiency between fractions. Such biases can distort fractionation profiles and be misinterpreted as PTM-dependent localisation changes. Similar effects were highlighted in phospho-thermal proteome profiling (PhosphoTPP), where separate enrichment of fractions introduced artefactual differences in phosphopeptide abundance unrelated to biological effects (Potel *et al*., 2021). Pooling fractions prior to enrichment therefore provides a more robust strategy for detecting genuine phosphorylation-dependent localisation differences. This approach also avoids two rounds of TMT labelling.

**Figure 5:**
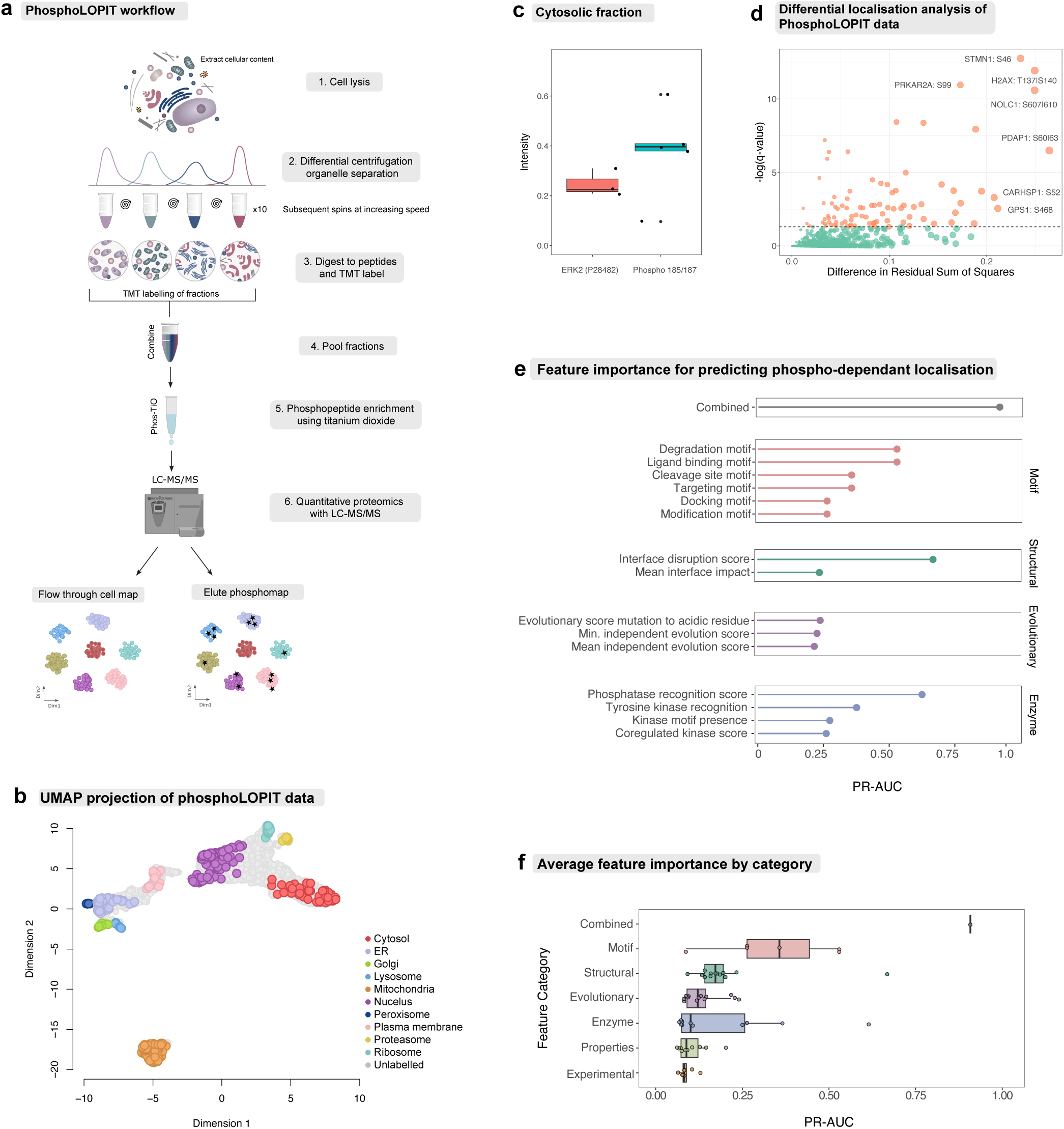
PhosphoLOPIT analysis using SANDLE. A) Schematic of the PhosphoLOPIT workflow (see methods) B) A t-SNE dimen-sionality reduction of the PhosphoLOPIT data where each pointer is a protein. Marker proteins are coloured. C) Cytosolic relative abundance of phosphorylated and unmodified peptides from ERK2. D) Differential localisation volcano plot showing the difference in residual sum of squares against the negative log of q-values (adjusted p-values). Dashed line indicates an FDR threshold of 0.1. Phosphosites above this line are coloured in orange; otherwise, they are coloured in green. E) Phosphoscore analysis showing highest scoring features using area under the curve for precision-recall analysis (PR-AUC). Quantities are colour coded according to their feature-type. F) Mean PR-AUC and standard error for feature categories. Number in each category indicated in the plot.

Focusing on the PhosphoLOPIT experiments, we made minor adaptations to the LOPIT-DC protocol and confirmed high-resolution of all organelles (Fig. 5 B, Supplementary Fig. 4). We found that Phosphoproteins were distributed throughout the subcellular environment with a bias towards nuclear and cytosolic localisations (Supplementary Fig. 5). One example was the activity-regulating phosphosites overlapping residues 185 and 187 of ERK2, an essential component of the MAP kinase signal transduction pathway. While the substrates of ERK2 are distributed throughout the cell and so ERK2 is multi-localised (Costa *et al*., 2006), we found that phosphorylation of the TXY activation motif (Thr-185 and Tyr-187) coincided with cytosolic enrichment of these phosphoproteins (Fig. 5 C). This is consistent with phosphorylation-dependent regulation of ERK2 signalling, although these data do not distinguish whether phosphorylation is a driver or consequence of localisation.

We applied SANDLE to identify modification-dependent localisation patterns by analysing proteins with matched modified and unmodified peptides, yielding 1,397 valid peptide pairs for testing. At a FDR of 0.05, we identified 87 proteins exhibiting phosphorylation-dependent localisation. Fig. 5 D highlights several proteins with dramatic shifts in subcellular distribution between their unmodified and phosphorylated states. One example is Selenon, which primarily localises to the endoplasmic reticulum (ER) in its unmodified form (Petit *et al*., 2003). Upon phosphorylation at serine-18, Selenon displays a distinct subcellular distribution, suggesting that this modification prevents signal peptide cleavage and enables Selenon functions outside the ER (Supplementary Fig. 6).

To investigate the molecular features associated with phosphorylation-dependent localisation, we made use of a previously developed functional phosphosite score that integrates proteomic, structural, regulatory and evolutionary features via machine learning (Ochoa *et al*., 2020). Our aim here was to assess which classes of features are informative about localisation changes, rather than to construct or evaluate a predictive model: the score and its constituent features are used purely as a vocabulary of biologically interpretable annotations. Treating SANDLE-identified phosphorylation-dependent localisation events as the positive class (FDR < 0.1), we computed the precision–recall area under the curve (PR-AUC) for the combined score and for each underlying feature. PR-AUC summarises how well a score separates phosphosites with localisation changes from those without, with values closer to one indicating stronger association; we interpret it here as a measure of feature relevance rather than out-of-sample predictive performance. The combined score achieved a PR-AUC of 0.908, and we next examined which individual features and feature categories drove this signal.

To understand this performance, we evaluated the predictive power of 81 individual feature-level predictors ((Ochoa *et al*., 2020)). Interface disruption scores (Guerois *et al*., 2002) (PR-AUC = 0.668) and phosphatase recognition sites (Blom *et al*., 2004) (PR-AUC = 0.615) were the strongest predictors (see Fig. 5E and methods). Tyrosine kinase recognition was also strongly predictive (PR-AUC = 0.37), consistent with the well-established role of phosphotyrosine in driving SH2/PTB-mediated recruitment and spatially regulated signalling (Pawson, 2004) and paralleling our recovery of phosphotyrosine adapters (GRB2, GAB1, SHC1) in the EGFR inhibition case study. These scores suggest that phosphorylation-associated localisation changes are consistent with modulation of protein-protein interactions, potentially by altering binding interfaces or causing conformational changes that expose or hide localisation signals. The phosphatase recognition score indicates these localisation-regulating sites are subject to dynamic spatio-temporal regulation, where the ability to be rapidly de-phosphorylated may be crucial for spatial control. Potential confounders such as protein disorder, protein abundance (PaxDB) (Wang *et al*., 2015), spectral counts, protein length and literature coverage (PubMed reference counts) showed minimal predictive power (PR-AUC < 0.11) indicating results are not driven by annotation bias. Rather this analysis suggests phosphorylation-dependent localisation is driven by local protein features. At the category level (see Fig. 5 F), linear motif annotations (Kumar *et al*., 2020) showed the strongest mean predictive power (PR-AUC = 0.341) followed by structural features (PR-AUC = 0.204) and enzyme recognition sites (PR-AUC = 0.188). The stronger performance of the combined score suggests multiple molecular mechanisms contribute to phosphorylation-dependent localisation changes, underscoring the complex nature of this regulatory mechanism.

To gain mechanistic insight into phospho-dependent localisation, we next examined which functional feature classes best explained individual sites. While the global analysis identified several predictive feature types, this does not indicate whether localisation is typically driven by a single dominant mechanism or by multiple contributing factors. We therefore grouped annotations into functional categories and, for each phosphosite, compared the strongest and second strongest category scores. Structural/interface annotations, kinase and phosphatase recognition features, and protein property annotations most frequently provided the strongest signal. Motif-based annotations were rarely dominant, but often showed moderate support alongside other feature classes. The strongest and second strongest category scores were frequently similar in magnitude (median ratio = 1.34), indicating that most phospho-dependent localisation events are supported by multiple feature classes rather than a single dominant mechanism (Supplementary. Fig. 7 and 8). Together, these results suggest that phosphorylation-dependent localisation typically reflects coordinated effects on protein interactions, targeting determinants, and regulatory recognition.

### Co-analysing re-localisation of transcriptome and proteome with SANDLE

SANDLE is equally applicable to spatial transcriptomics datasets generated by subcellular fractionation. Recently, phase-separation coupled fractionation enabled simultaneous tracking of RNA (LoRNA) and protein (LOPIT) localisation dynamics, presenting new analytical challenges (Fig. 6A) (Villanueva *et al*., 2024). To evaluate SANDLE in this context, we analysed U-2 OS cells treated with thapsigargin (Tg) for 1 h, which activates the unfolded protein response (UPR) (Villanueva *et al*., 2024). This perturbation induces global translational repression and promotes re-localisation of both proteins and transcripts into stress granule-associated fractions (Sehgal *et al*., 2017).

**Figure 6:**
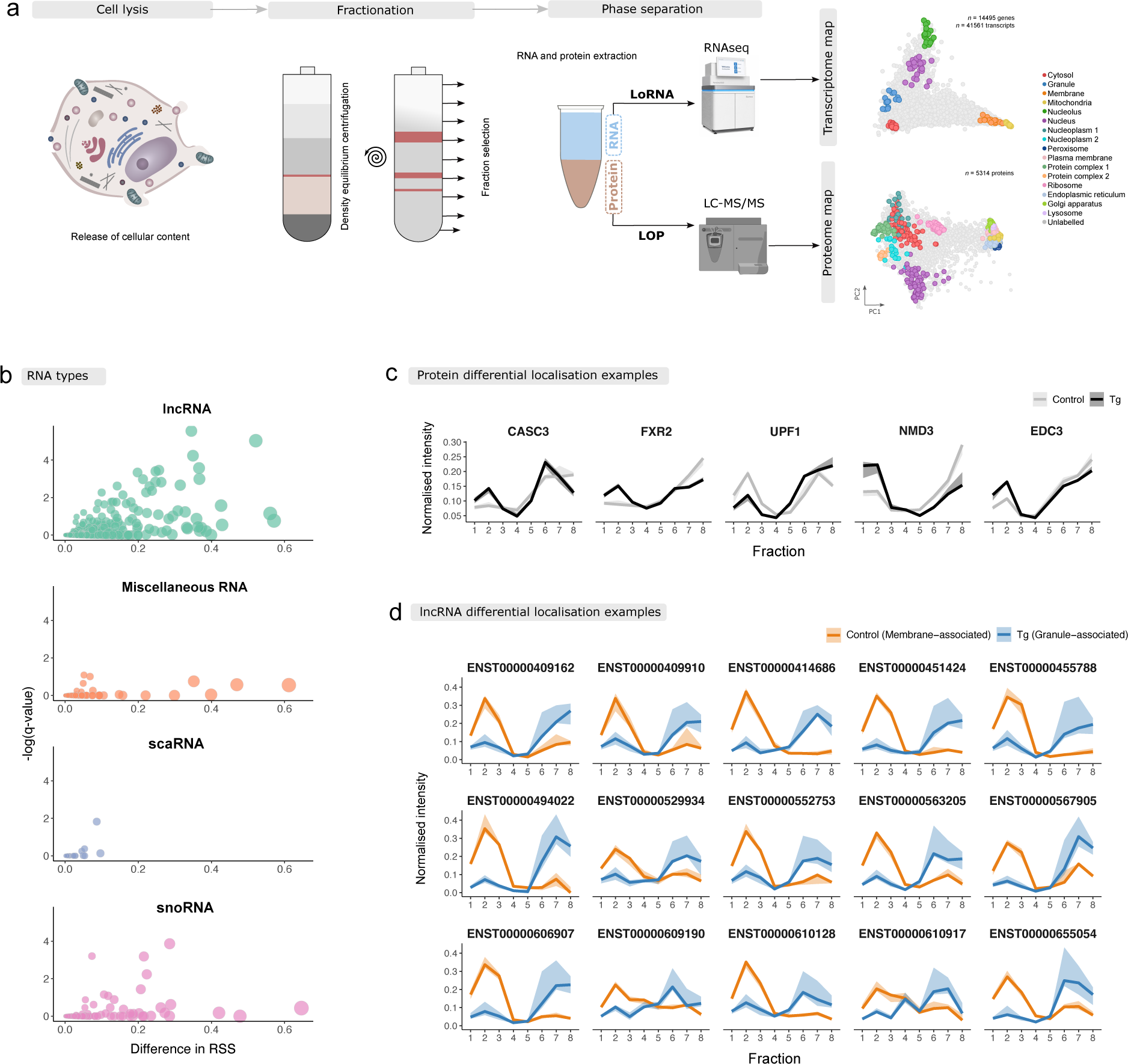
SANDLE analysis of multi-omic subcellular data. A) The multi-omic subcellular workflow combining both localisation of RNA and localisation of protein by isobaric tagging. B) Volcano plots faceted by RNA type. The difference in residual sum of squares (localisation change effect size) is plotted against the -log(q-value). The pointers are also scaled by the effect size and coloured according to RNA type. C) Example differentially localised proteins from the protein-level data determined using SANDLE D) The lncRNAs with localisation changes from membrane to cytosolic granules.

We first analysed protein re-localisation following thapsigargin treatment using the following thresholds: FDR < 0.05, DL probability > 0.5, ΔRSS > 0.5. SANDLE identified 91 proteins exhibiting spatial reorganisation, with enrichment of RNA-associated factors (18% of relocalised proteins). Notably, FXR2, UPF1, CASC3 and NMD3, all involved in RNA metabolism, shifted towards cytosol-light fractions associated with stress granules, consistent with the original LoRNA study (Villanueva *et al*., 2024). We also observed re-localisation of EDC3, a P-body assembly factor and decapping activator, supporting reorganisation of cytoplasmic RNA granules in response to Tg treatment (Fig. 6B). These changes suggest formation or re-organisation of stress-granules in response to Tg treatment. Several relocalised proteins participate in mRNA surveillance and post-transcriptional regulation, including UPF1 and CASC3. During cellular stress, transcripts are sequestered into RNA granules, where they are protected from degradation and translational processing. This sequestration is thought to preserve transcripts during stress recovery and prevent production of misfolded proteins (Goetz and Wilkinson, 2017; Hug *et al*., 2016). The observed re-localisation of RNA regulatory factors is therefore consistent with stress granule assembly following ER stress.

SANDLE also identified the re-localisation of calcium/calmodulin-dependent protein kinase type II subunits gamma and delta (CAMK2G and CAMK2D) (see Supplementary Fig. 9). This represents an interesting calcium signalling response to thapsigargin treatment. As thapsigargin inhibits SERCA pumps, resulting in elevated cytosolic calcium levels, the spatial redistribution of these calcium-responsive kinases could reflect functional adaptation during ER stress (Timmins *et al*., 2009). The observed re-localisation pattern of CAMK2G/D may reflect parallel or coordinated cellular response to RNA metabolism changes caused by thapsigargin-induced stress. A causal link requires further study but represents an interesting hypothesis that CAMK2 mediated singalling may contribute to RNA metabolism during UPR.

We next leveraged SANDLE to analyse the subcellular transcriptomics data comprising 41, 561 transcripts. Despite this dataset’s considerable size, analysis was completed within minutes, demonstrating SANDLE’s computational efficiency for large-scale spatial transcriptomics. First, we examined whether any of the proteins we previously mentioned also exhibited transcript-level re-localisation, finding no evidence for coordinated RNA-protein spatial dynamics. Given the stress granule associations observed in our proteomics data, we sought to understand lncRNA’s localisation changes upon thapsigargin treatment. Curiously, SANDLE identified 29 differentially localised lncRNAs (Fig. 6 C), with 17 exhibiting a distinct shift from membrane-enriched fractions (peaking in fraction 2) to granule-associated fractions (peaking in fractions 6-7) see Fig. 6 D. This membrane-to-granule re-localisation pattern may represent a conserved regulatory mechanism that modulates gene expression during ER stress. Among these lncRNAs was PINK1-AS, an antisense transcript that regulates expression of the mitochondrial kinase PINK1 (Scheele *et al*., 2007; Chiba *et al*., 2009). PINK1 is a key regulator of mitochondrial quality control and is implicated in Parkinson’s disease, suggesting that stress-induced re-localisation of PINK1-AS may reflect regulation of mitochondrial stress pathways.

The coordinated re-localisation of multiple lncRNAs suggests a structured cellular response rather than non-specific aggregation. RNA granules may act as transient storage compartments where transcripts are protected from degradation and translational processing during stress. Furthermore, to determine whether relocalising lncRNAs represent a distinct subset, we compared their sequence features to the broader population of detected lncRNAs. While GC content, G-quadruplex motif abundance and transcript length showed no detectable association, we note that the modest number of relocalising lncRNAs limits power to identify weaker effects (Supplementary Fig. 10). In contrast, translation-associated features showed the strongest and most consistent signal (Supplementary Fig. 10). Relocalising lncRNAs exhibited increased 5’ AUG density but tended to lack long open reading frames, suggesting that these transcripts are recognised by the translational machinery but are unlikely to support productive translation (Supplementary Fig. 10).

These observations is consistent with a model in which RNA re-localisation during stress is not a passive consequence of global translational shutdown, but instead reflects selective partitioning based on engagement with the translation machinery. In this framework, stress granules act as sites of translation triage, preferentially sequestering transcripts that initiate but fail to sustain productive translation. This provides a mechanistic explanation for the behaviour of this lncRNA subset and highlights how SANDLE can move beyond identifying re-localisation events to uncovering sequence-level determinants of spatial organisation in the cell. Together, these results demonstrate SANDLE’s versatility for analysing complex multi-omic subcellular fractionation datasets, enabling unified detection of protein and RNA re-localisation events.

## Discussion

We have presented SANDLE, a fast, robust, and modular framework for analysing subcellular omic data. By integrating marker-free and marker-dependent approaches, SANDLE enables the analysis of diverse differential localisation experimental designs, including multi-omics subcellular data. Our comparative analyses demonstrate that SANDLE outperforms existing tools such as TransGCN in both accuracy and computational efficiency, being up to 100-fold faster. Unlike other methods that show inflated false discovery rates at commonly used thresholds, SANDLE maintains proper error control even with complex experimental designs. More generally, our results suggest that robust differential localisation is best viewed as a problem requiring independent evidence from both localisation assignment and profile perturbation. This principle is independent of any particular statistical model and may extend naturally to future spatial omics technologies.

A practical consequence of SANDLE is that we can give power estimates for spatial proteomics experiments. With the field-standard three-replicate design, our simulations indicate that approximately one third of true differential localisation events remain undetected at FDR < 0.05. This a substantial under-counting that has been masked by the inflated false positive rates of existing methods, where apparent sensitive detection partly reflects miscalibrated FDR control. Each additional replicate contributes 0.012 to AUC and each additional fraction contributes 0.007 across the regimes we tested, providing quantitative guidance for design trade-offs. Together with SANDLE’s 100-fold speedup, these results have direct implications for experimental scale: clinical cohorts, ligand titrations, and longitudinal designs that were previously gated by either statistical or computational cost are now accessible. Re-analysis of published datasets is also likely to recover differential localisation events missed by earlier methods, particularly in studies with limited replication.

SANDLE’s utility extends across various biological contexts. In well-characterized systems, it successfully recovered known differential localisations associated with signalling responses (Insulin in adipocytes), drug treatments (EGFR inhibitor Gefitinib), and cell-type differences (CHO-K1 versus MPC-11). While these findings validate SANDLE’s performance, its greater value lies in uncovering novel insights in challenging scenarios.

In our parasitology example, SANDLE uncovered previously unreported differential localisations between blood stream and procyclic forms of *Trypanasoma brucei*, as well across species between *T. brucei* and *T. congolense*. Furthermore, a known glycosomal marker in *T. brucei* appeared to have a different localisation in *T. congolense*. Curiously, other proteins with correlated profiles also appear to have altered localisation in a concerted manner suggesting fundamental difference in compartmentalisation between the species and therefore potential changes in metabolic requirements of the two species. These findings demonstrate SANDLE’s capacity to generate new biological hypotheses in complex comparative systems. Application of SANDLE to PhosphoLOPIT, a novel subcellular phospho-proteomics dataset, identified numerous proteins with phosphorylation-dependent localisation. Through analysis of 81 feature-level predictors, we found that interface disruption scores and phosphatase recognition sites has the strongest feature association with phosphorylation-dependent localisation. This is consistent with a mechanistic link between phosphorylation, protein-protein interaction interfaces, and spatial regulation, whereby modification of interface residues drives conformational changes that alter subcellular targeting. While these findings are compelling, we acknowledge limitations in our ability to distinguish direct phosphorylation effects from downstream consequences; future targeted studies will be required to establish causality.

When applied to multi-omic subcellular data, SANDLE revealed complex post-transcriptional regulation in response to Thapsigargin treatment. We observed re-localisation of many proteins, including EDC3, a P-body assembly factor that showed differential localisation. Concurrently, 17 lncRNAs exhibited enhanced granule association following Thapsigargin treatment. This parallel behaviour of proteins and RNAs suggests a coordinated stress response mechanism involving lncRNAs, an observation that would be missed by protein-only analyses. A limitation of our approach is the inability to determine whether protein re-localisation drives RNA re-distribution or vice versa; complementary approaches will be needed to resolve this question.

Future development will focus on extending SANDLE to analyse temporal changes of protein localisation and incorporating clinical covariates. We are also exploring integration with protein structural information to better predict how modifications affect localisation. The SANDLE framework is available as an open-source R package within the BANDLE Bioconductor package.

In summary, SANDLE represents a significant advance in subcellular omics analysis. Its application has already revealed novel insights into compartment-specific regulation, species-specific differences in organelle composition, and stress-induced reorganisation of the protein-RNA interactome. By enabling more analysis of the dynamic spatial organization of cellular components, SANDLE provides a valuable tool for understanding how subcellular relocalisation contributes to cellular function and adaptation. Existing differential localisation methods have largely sought increasingly sophisticated probabilistic models of localisation. SANDLE instead takes a modular view, recognising that localisation assignment and profile perturbation are distinct statistical questions that can be addressed using complementary methodologies.

## Methods

### SANDLE statistical framework

SANDLE combines a marker-dependent generative model of subcellular niches with a marker-free regression-based test for changes in fractionation profiles.

### Marker-dependent modelling of subcellular niches

For each experiment, proteins with known subcellular localisation (markers) were used to define *K* subcellular niches. Let **x***_i_* ∈ R*^d^* denote the fractionation profile of protein *i* across *d* fractions. For each niche *k* ∈ {1*, …, K*}, we modelled marker profiles using a multivariate *t*-distribution:

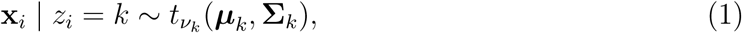

where ***µ****_k_* is the niche-specific mean profile, **Σ***_k_* is the covariance matrix, and *ν_k_* is the degrees of freedom.

Robust estimates of ***µ****_k_* and **Σ***_k_* were obtained using minimum covariance determinant estimation, with fallback to empirical estimates where necessary due to numerical errors. To ensure positive-definite covariance estimates in high-dimensional settings, graphical lasso regularisation was applied:

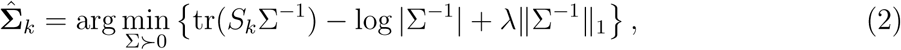

where *S_k_* is the empirical covariance matrix and *λ* is a regularisation parameter. Degrees of freedom *ν_k_* were estimated by maximum likelihood.

### Posterior allocation probabilities

For each protein *i*, posterior probabilities of belonging to each niche were computed under a finite mixture model:

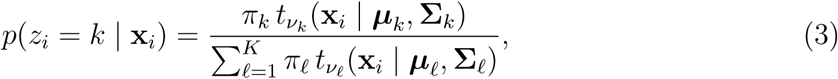

where *π_k_* are mixture weights estimated from marker frequencies. These probabilities define a soft assignment of each protein to subcellular niches.

### Marker-free test for differential localisation

Independently of marker annotations, differential localisation was assessed using a regression-based test on fractionation profiles. For each protein *i*, replicate *r*, condition *j* ∈ {1*, …, J*} and fraction *k* ∈ {1*, …, d*}, we model the abundance *y_ijkr_* as

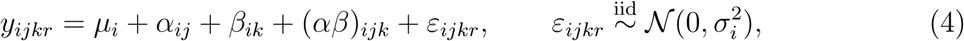

where *µ_i_* is a protein-specific intercept, *α_ij_* is a main effect of condition, *β_ik_* is a main effect of fraction, and (*αβ*)*_ijk_* is the condition-by-fraction interaction. Sum-to-zero contrasts are applied to *α_ij_*, *β_ik_* and (*αβ*)*_ijk_* for identifiability.

Differential localisation manifests as a non-zero interaction term: a change in overall abundance between conditions affects *α_ij_* but leaves the fractionation profile unchanged, whereas a change in subcellular distribution alters the *shape* of the profile across fractions and is captured by (*αβ*)*_ijk_*. We therefore test

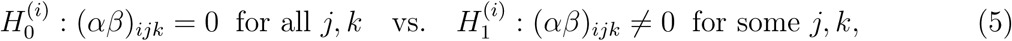

using an *F*-test comparing the full model against the additive (no-interaction) reduced model. The resulting per-protein *p*-values *p_i_* are adjusted across proteins using the Benjamini–Hochberg procedure to obtain FDR*_i_*.

The associated effect size is the difference in residual sum of squares between the reduced and full models,

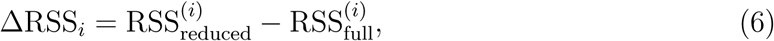

which quantifies the magnitude of the condition-dependent change in fractionation profile and provides an interpretable analogue of log fold change for ranking and visualisation.

### Differential localisation probability

To quantify the probability that a protein changes subcellular localisation between conditions, we sampled niche assignments from the posterior distributions. For each protein *i*, we drew samples

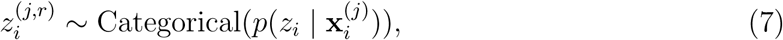

for replicate *r* in condition *j*.

For two conditions, the differential localisation probability was defined as

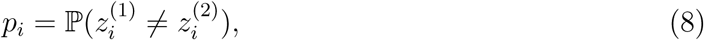

estimated as the fraction of sampled allocations that differ between conditions. Both paired and unpaired estimators were considered depending on experimental design.

### Combined decision rule

SANDLE combines the marker-free statistical test and the marker-dependent probability into a joint score:

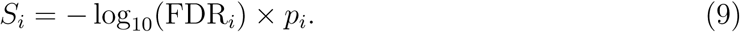

This quantity is used for AUC ranking only not for calling due to the different scales and lack of independence of the two measurements. In addition to ranking by *S_i_*, proteins were called differentially localised using joint thresholds on both quantities:

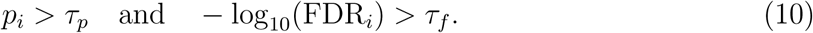

This formulation separates statistical evidence for a change in profile from the probability of a change in subcellular assignment.

### Final allocation across replicates

For visualisation and interpretation, final niche assignments were obtained by combining posterior probabilities across replicates:

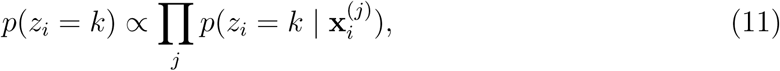

followed by normalisation and maximum a posteriori assignment.

### Datasets and Preprocessing

All datasets apart from the PhosphoLOPIT data (described below) were taken as supplied in the original manuscripts. No additional imputation was performed and marker definitions were not changed. Data was analysed at the Protein level (proteomics datasets) or transcript level (transcriptomics datasets). Profiles were median-centred across fractions prior to marker-free testing to remove global intensity effects.

### Simulation benchmarking

SANDLE was benchmarked using 60 simulated LOPIT experiments with known ground-truth differential localisation status. Simulations were generated from the Tan (2009) spatial proteomics dataset and consisted of two experimental conditions with three replicates per condition. In each simulation, a fixed set of 30 proteins was assigned as differentially localised. Simulation scenarios varied nuisance structure including batch effects, fraction permutation and bootstrap strategy, allowing performance to be assessed across a range of realistic experimental artefacts as previously described (Crook *et al*., 2022).

For each simulated dataset, SANDLE was applied to the six replicate-level datasets. Differential localisation probabilities were computed using the unpaired design:

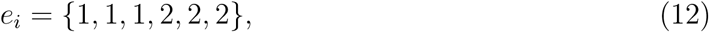

where *e_i_* denotes the experimental condition for each replicate. In parallel, the marker-free regression test was applied to the same simulated profiles. Marker proteins were assigned a differential localisation probability of zero before performance evaluation, to avoid scoring training markers as candidate differential-localisation events. The marker-dependent probability, marker-free adjusted significance and ground-truth status were then combined into a single results table for each simulation.

Performance was assessed using area under the receiver operating characteristic curve (AUC). Three SANDLE scores were compared: the marker-free score, defined as − log_10_(FDR); the marker-dependent differential localisation probability; and a combined score given by

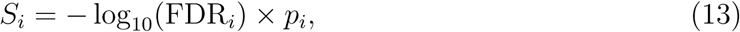

where *p_i_* is the SANDLE differential localisation probability for protein *i*. SANDLE was compared with BANDLE and TransGCN on the same 60 simulated datasets. For BANDLE, posterior differential localisation probabilities were extracted from the fitted model and used directly as the ranking score. For TransGCN, protein translocation probabilities and estimated FDR values were read from the TransGCN output files. AUCs were computed using the known set of simulated differentially localised proteins as positives.

In addition to ranking performance, we assessed performance at recommended operating thresholds. For SANDLE, proteins were called differentially localised if they satisfied both

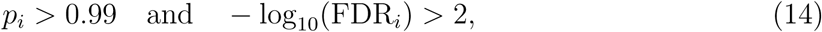

corresponding to a differential localisation probability threshold of 0.99 and FDR *<* 0.01. A relaxed SANDLE threshold was also evaluated:

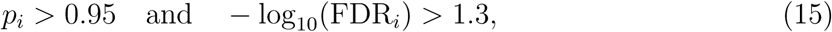

corresponding approximately to FDR *<* 0.05. BANDLE calls were made using posterior differential localisation probability *>* 0.99, while TransGCN calls were made using estimated FDR *<* 0.01. For each method and simulation, we recorded true positives, false positives, false negatives, sensitivity and false positive rate.

Empirical false discovery rates were estimated by comparing the called proteins against the known ground truth. For a given threshold, the observed false discovery rate was computed as

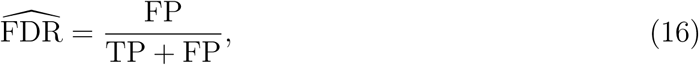

where FP and TP denote the number of false-positive and true-positive calls, respectively. Estimated and observed FDR values were compared to assess calibration.

Runtime was recorded for each SANDLE simulation using wall-clock time around the SANDLE and marker-free testing steps. BANDLE runtimes were taken from stored benchmarking output. TransGCN runtimes were estimated from output-file modification times across sequentially completed simulations. Runtime values were converted to minutes and compared across methods.

### Experimental design and power simulations

To assess how experimental design affects SANDLE performance, we analysed simulated LOPIT datasets with varying numbers of subcellular fractions and biological replicates. Simulated datasets were generated with between four and nine fractions and between two and five replicates per condition. Each dataset contained two experimental conditions and a known ground-truth set of 30 differentially localised proteins.

For each combination of fraction number and replicate number, SANDLE was applied to the replicate-level datasets. Differential localisation probabilities were computed using an unpaired design:

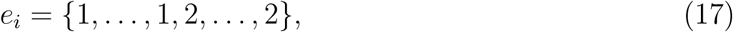

where the number of repeated entries for each condition corresponded to the number of replicates. The marker-free regression test was then applied to the same simulated profiles. Marker proteins were assigned differential localisation probability zero before performance evaluation.

For each simulation setting, proteins were called differentially localised using the relaxed SANDLE thresholds

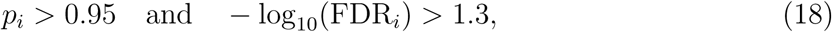

where *p_i_* is the differential localisation probability for protein *i*. This corresponds approximately to FDR *<* 0.05. For each design, we recorded the number of true positives, false positives, false negatives, sensitivity and false positive rate.

To compare marker-free, marker-dependent and combined scoring across experimental designs, we computed receiver operating characteristic AUCs for three ranking scores:

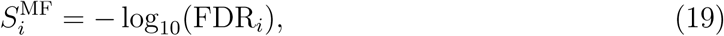

and

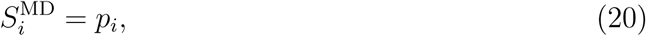

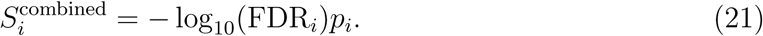

Here, *S^MF^_t_* is the marker-free score, *S^MD^_t_* is the marker-dependent score, and *S^combined^_t_* is the combined SANDLE score.

Finally, to quantify the relative contribution of experimental design parameters to detection performance, we fitted a linear model:

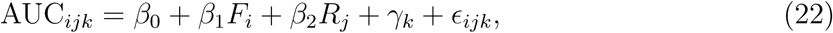

where *F_i_* is the number of fractions, *R_j_* is the number of replicates, and *γ_k_* denotes the scoring method. This model was used to estimate the marginal effect of adding fractions or replicates on AUC.

### Cell Culture

The U-2 OS human bone osteosarcoma cell line (generous gift from Prof. Emma Lundberg, Science for Life Laboratory, School of Biotechnology, KTH Royal Institute of Technology, SE-171 21 Stockholm, Sweden.) was cultured and incubated at 37 degrees in McCoy’s 5A (Gibco™; 16600082) supplemented with 10% FBS in humidified conditions with 5% CO2, without antibiotics. The cell line was tested to confirm the absence of Mycoplasma.

### Sample preparation

Cells were harvested at 90% confluence by trypsinisation, washed with PBS three times and resuspended in lysis buffer consisting of 0.25M sucrose, 10mM HEPES pH 7.4, 2mM EDTA, 2 mM magnesium acetate, protease inhibitor (Roche) and phosphatase inhibitor (PhosStop Roche). Cell lysis was performed with ball bearing homogenizer (isobiotec) where each 1.5 ml of cells was passed through homogeniser chamber 25 times with 12*µ*m ball-bearing clearance size. The lysate was collected and differential centrifugation was performed for subcellular fractionation.

### PhosphoLOPIT subcellular fraction

To obtain subcellular fractions for phosphoproteome enrichment cell lysate was fractionated using differential centrifugation. For each biological replicate, cell lysate was separated into ten subcellular fractions: 100g (10 min), 500g (5 min), 1000g (10 min), 3000g (10 min), 5000g (10 min), 9000g (15 min), 15,000g (15 min), 30,000g (20 min), and 120,000g (45 min). Centrifugation steps up to 30,000*g* were performed with Eppendorf 5424R benchtop centrifuge; the last two centrifugation steps were carried out using a Beckmann Optima MAX-XP Ultracentrifuge. Pellets from nine fractions and the supernatant from the last fraction were collected and stored in −20 degrees Celsius. Collected pellets were solubilised using a membrane solubilisation buffer which contained 8M Urea, 0.2% (wt/vol) SDS, 50mM HEPES (pH 8.5), protease and phosphatase inhibitors. Nuclease enzyme was added to the first three centrifuged fractions to break down interfering DNA before protein concentration quantitation was performed with BCA kit (Thermo Fisher Scientific) according to manufacturer instructions. After quantitation, all fractions were precipitated with ethanol/acetone to remove urea, protease and phosphatase inhibitors that could interfere with protein digestion. Briefly 400 *µ*l of ice-cold ethanol and 400 *µ*l of ice-cold acetone to each 100 *µ*l of sample, vortexed and incubated at −20 degrees overnight. Following day, the samples were centrifuged at 16000*g* for 15 minutes at 4 degrees and the supernatant discarded.

### Protein digest and peptide labelling

Precipitated subcellular fractions were solubilised in 100mM TEAB, reduced with 15mM dithiothreitol (DTT) for 1 hour at 37 degrees followed by alkylation with 55 mM iodoacetamide (IAA) for 1 hour at room temperature in the dark. Two step digestion was performed, first with 3*µ*g of trypsin (Promega) overnight at 37 degrees followed by further digestion with 1*µ*g of trypsin for four hours. Fractions were then acidified with TFA (0.1% final concentration), centrifuged to pellet any debris and peptide supernatant desalted. For peptide clean-up peptide desalting spin columns were used (Pierce) according to manufacturer’s guideline. Briefly, spin columns were conditioned, fractions loaded, cleaned-up and eluted with 50% acetonitrile 50% HPLC grade water 0.1% trifluoroacetic acid (TFA). Collected peptides from the ten subcellular fractions were quantified with Qubit (manufacturers details) and equivalent amounts of peptides from each fraction were taken and dried before TMT labelling (Thermo Fisher Scientific). For each biological replicate, between 80-100*µ*g of peptides were labelled per subcellular fraction, resulting in each TMT multiplex being between 800*µ*g - 1000*µ*g. Fractions were labelled according to manufacturer’s protocol. In short each TMT tag (0.8mg) was resuspended in acetonitrile and incubated for 1 hour with a desalted sample digest that was resuspended in 100mM TEAB before TMT labelling. Each TMT reaction was quenched with 5% hydroxylamine for 15 mins and samples subsequently combined in equal proportions (multiplex). 10% of each multiplex was taken for total proteome analysis and 90% of the multiplex was taken for phosphoproteome enrichment.

### Phosphopeptide enrichment

For each biological replicate, 90% of the multiplex was dried down and stored at −20 degrees before phosphopeptide enrichment. Phosphopeptide enrichment was performed using TiO_2_ enrichment and SIMAC (sequential elution from IMAC) methodology, a strategy separately enriching monophosphorylated and multi-phosphorylated peptides. Briefly, each lyophilized multiplexed sample was resuspended in 1 M glycolic acid in 80% acetonitrile 5% TFA and incubated with 0.6 mg of TiO2 beads per 100 *µ*g of peptide sample under vigorous shaking for 15 min. Beads were spun down and the supernatant was transferred to a new eppendorf tube to which fresh beads were added, half of the original amount of TiO2, sample again was incubated under vigorous shaking for 15 mins. This process was repeated two more times. The TiO2 beads from the three rounds of enrichment were combined and washed first with 80% acetonitrile 0.1% TFA and then with 10% acetonitrile 0.1% TFA to remove non-phosphorylated peptides bound to TiO_2_. Phosphopeptides were then eluted from the beads with a fresh ammonia elution buffer (NH4OH - 0.28%), pH 11, before being dried down. This enriched sample was solubilised in 150 *µ*l SIMAC loading buffer (50% acetonitrile 0.2% TFA). For each of these enriched samples, 80 *µ*l of iron-coated PHOS-selectTM Iron Affinity gel (metal chelate (IMAC) beads) (Sigma) was used. The beads were washed twice with 1 mL SIMAC loading buffer, then combined with 150 *µ*l sample solution and incubated under vigorous shaking for 30 min at room temperature. After incubation, the IMAC beads were packed in the constricted end of a 200 *µ*l GELoader tip by air pressure. The IMAC flow-through was collected in a new eppendorf. Packed IMAC beads were then washed twice with 50 *µ*l of SIMAC loading buffer and with 70 *µ*l of 20% acetonitrile 1% TFA. The washes were combined with the flow-through, this flowthrough was enriched in monophosphorylated peptides. Multi-phosphopeptides were then eluted from the IMAC column with ammonia solution to a new eppendorf tube. The multi-phosphorylated peptides were acidified with 10 *µ*l 100% formic acid and purified by Oligo R3 reverse-phase micro-columns. Monophosphorylated fraction was dried down, resolubilised in 70% acetonitrile 2% TFA and further enriched with two rounds of TiO_2_. After incubation, the beads were washed with 50% acetonitrile 0.1% TFA, eluted with 100 *µ*l of fresh ammonia solution and dried down. Monophosphorylated sample was then pre-fractionated with High pH Reversed-phase Peptide Fractionation Kit from Pierce according to the manufacturer’s instruction. Briefly, the fractionation column was conditioned, the sample loaded and fractions collected with increasing percentage of acetonitrile resulting in eight monophosphorylated fractions. The resulting eight monophosphorylated fractions and one multi-phosphorylated fractions were dried fully, solubilised in 20 *µ*l 0.1% formic acid. Half of the sample was analysed by LC-MS/MS as described below.

### Off-line UPLC fractionation

For total proteome analysis, 10% of the TMT10-plex sample was resuspended in 100 *µ*l of 20mM ammonium formate in HPLC grade water (buffer A) and pre-fractionated using basic pH reverse-phase chromatography, Acquity UPLC system from Waters equipped with an autosampler, a binary solvent manager and a diode array detector. The total proteome sample was loaded to Acquity UPLC BEH C18 column (2.1-mm i.d. × 150-mm; 1.7-*µ*m particle size) where following buffers for gradient elution were used; 20 mM ammonium formate for buffer A and 20 mM ammonium formate 80% acetonitrile pH 10 for buffer B. A flow rate of 0.244 ml min-1 was maintained. The percentage of buffer B was varied according to the following programme: 5% for 10 min, 5 to 75% over 50 mins, a ramp to 100% over 2 mins followed by 5.5 mins at 100%, switching to 5%, and equilibration for 10 min. Between 36 and 38 fractions were collected along the elution profile of the peptides (approximately from minute 22 to 60 of the program) and concatenated (sample 1 with sample 16 etc…) into 15 fractions which eluted at different time points during the gradient. These fractions were dried down and solubilised in 0.1% formic acid where approximately 1.5*µ*g of peptides were loaded per mass spectrometry run.

### LC-SPS-MS/MS/MS

Both total and phospho proteome samples were analysed in an Orbitrap Fusion Lumos Tribrid mass spectrometer (Thermo Fisher Scientific), coupled to a nanoUPLC (Dionex Ultimate 3000 UHPLC). Peptides were loaded onto a trap-column (Thermo Fisher Scientific PepMap 100 C18, 5 *µ*m particle size, 100A pore size, 300 *µ*m i.d. x 5mm length) and separated by C18 reverse-phase chromatography at a flow rate of 300 nL/min and a Thermo Scientific reverse-phase nano Easy-spray column (Thermo Scientific PepMap C18, 2*µ*m particle size, 100A pore size, 75*µ*m i.d. x 50cm length). Analytical chromatography both for total and phospho proteomes consisted of Buffer A (HPLC water/ 0.1% formic acid) and Buffer B (80% acetonitrile/ 0.1% formic acid). 0-3 min at 2% buffer B, 3-93 min linear gradient 2% to 40% buffer B, 93-100 min linear gradient 40% to 90% buffer B, 100-104 min at 90% buffer B, 104-105 min linear gradient 90% to 2% buffer B and 105-120 min at 2% buffer B. All samples were acquired in a positive ion mode, 120 min run and applying data dependent acquisition using synchronous precursor selection MS3 (SPS-MS3). Phosphoproteome samples utilised multiscan activation method to trigger fragmentation of parent ion and neutral loss. For both total and phospho proteomes peptide ions were measured in an Orbitrap mass analyzer, set at a resolution of 120,000 and were scanned between m/z 380-1500 Da. Data dependent MS/MS scans (3 second duty cycle time) were employed to automatically isolate and fragment precursor ions using Collisional-Induced Dissociation (CID) (Normalised Collision Energy of 35%). Only precursors with charge between 2 to 7 were selected for fragmentation and selected fragmented ions were dynamically excluded for 70 s. For total proteome MS2 fragments were measured with the Ion Trap analyser. AGC target was set to 10,000, maximum accumulation time to 50 ms and precursor isolation was performed by the quadrupole with 0.7 m/z transmission window. SPS was applied to co-select 10 fragment ions to undergo Higher energy Collisional-induced Dissociation (HCD) fragmentation with 65% normalized collision energy. SPS ions were all selected within the 400–1,200 m/z range and orbitrap resolution of 50,000. AGC targets and maximum accumulation times were set to 200,000 and 120 ms respectively. For phosphoproteome MS2 fragments were measured with the Orbitrap analyser with 30,000 resolution. AGC target of 50,000 and maximum accumulation time to 60 ms were employed. Multistage activation was enabled and neutral loss mass set to 97.9673. SPS was applied to co-select 10 fragment ions to undergo Higher energy Collisional-induced Dissociation (HCD) fragmentation with 65% normalized collision energy, orbitrap resolution of 60,000. SPS ions were all selected within the 400–1,200 m/z range and were set to preclude selection of the precursor ion and TMT ion series. AGC target and maximum accumulation time were set to 200,000 and 120 ms respectively.

### Peptide and protein identification

Raw data were viewed in Xcalibur v.3.0.63 (Thermo Scientific) and data processing was performed using Proteome Discoverer v.2.4 (Thermo Scientific). The raw files were searched against a Homo sapiens database (downloaded from uniport April 2018) and common contaminant database (cRAP) using Proteome Discoverer with SequestHF algorithm. The spectra identification was performed with the following parameters: MS accuracy 10 ppm; MS/MS accuracy 0.5 Da; up to two missed cleavage sites were allowed; carbamidomethylation of cysteine, TMT10plex tagging of lysine and peptide N-terminus were set to fixed modification; oxidation of methionine and deamidation of asparagine and glutamine were set to variable modifications. For phospho enriched fractions, phosphorylation of serine, threonine, and tyrosine residues were included as variable modifications. TMT reporter values were assessed through Proteome Discoverer v2.4 using the Most Confident Centroid method for peak integration and integration tolerance of 20 p.p.m. Reporter ion intensities were adjusted to correct for the isotopic impurities of the different TMT reagents (manufacturer specifications). Percolator node was used for false discovery rate estimation and only peptide identifications of high confidence (FDR<1%) were accepted. A minimum of one high confidence peptide per protein was accepted for identification using Proteome Discoverer. For phosphopeptide identifications, only high confidence phosphopeptides with phosphorylation site probability scores (ptmRS) above 75% were accepted.

### Data preprocessing

In brief, PSMs were removed if they matched a cRAP protein, did not have an assigned “master” protein, did not have quantification values, and had signal:noise ratio less than 5 or coisolation/interference above 50%. For the phospho multiplex, peptides without a phospho site with site probability scores above 0.75 were also removed. Up to three missing values were imputed using KNN (k = 10) with sum normalization beforehand and denormalization after imputation to ensure that nearest neighbors had similar profiles over the tags rather than a similar overall intensity. PSM intensities were then log center-median normalized before exponentiating back to the untransformed values. PSMs from the total multiplex were aggregated to peptide-level and protein-level intensities by summation of PSM-level tag intensities. Similarly, PSMs from the phosho-enriched multiplex were aggregated to succinyl-peptide level by summation of PSM-level tag intensities. Peptide- and protein-level quantifications were row-sum normalized.

### Functional annotation analysis of phosphorylation-dependent localisation

To investigate whether phosphorylation-dependent localisation changes were associated with known functional properties of phosphosites, we integrated SANDLE results with published phosphosite functional annotations. For each matched phosphopeptide–protein pair tested by SANDLE, we extracted the corresponding UniProt accession and phosphosite position. These identifiers were used to match sites to the functional phosphosite score described by Ochoa *et al*. (2020), together with the individual feature-level annotations used to construct that score.

Phosphosites were classified as showing phosphorylation-dependent localisation if they passed the SANDLE significance threshold used for this analysis. Specifically, sites, corresponding to an FDR threshold of 0.1. All remaining tested sites were assigned to the non-differential class. Sites without a matched functional annotation were excluded from score-based analyses.

We first asked whether the published functional phosphosite score was predictive of phosphorylation-dependent localisation. A weighted logistic regression model was fitted:

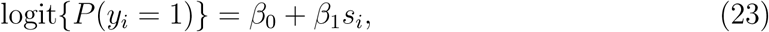

where *y_i_* indicates whether phosphosite *i* was classified as differentially localised and *s_i_* is the published functional score for that site. To reflect the higher confidence assigned to significant SANDLE hits, observations classified as differentially localised were assigned a larger weight than non-significant observations. Model predictions were used to assess discriminative performance using receiver operating characteristic and precision–recall analyses. In addition, the functional score itself was evaluated directly using precision–recall area under the curve (PR-AUC), with differentially localised sites treated as the positive class.

To determine which functional features were most informative, we next analysed individual phosphosite-level annotations. Feature annotations were matched to SANDLE results by UniProt accession and phosphosite position. For each feature, PR-AUC was computed after removing missing values:

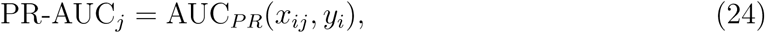

where *x_ij_* denotes the value of feature *j* for phosphosite *i*, and *y_i_* denotes differential localisation status. Features for which only one outcome class was represented after filtering were excluded.

Individual features were grouped into broad functional categories: structural/interface features, enzyme recognition features, short linear motif annotations, evolutionary features, experimental evidence, and general protein properties. Feature-level PR-AUC values were visualised individually and summarised within categories by computing the mean PR-AUC and its standard error:

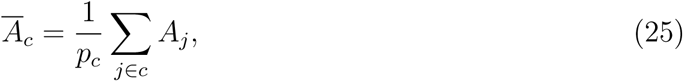

where *A_j_* is the PR-AUC for feature *j* and *p_c_* is the number of features in category *c*. This provided a category-level summary of which classes of phosphosite annotations were most associated with phosphorylation-dependent localisation. The full list of features and their descriptions is given as:

### Category-level analysis of phosphosite functional annotations

To gain mechanistic insight into phospho-dependent localisation, we summarised phosphosite annotations into functional feature categories. Individual phosphosite features were obtained from curated functional annotation datasets and grouped into broad categories, including structural/interface annotations, enzyme recognition features, short linear motif annotations, evolutionary conservation metrics, experimental evidence, and protein property annotations. Continuous features were standardised across sites, and binary or categorical annotations were encoded as numeric indicators where appropriate.

For each phosphosite, a category-level score was computed to summarise the strength of evidence for each functional class. Specifically, for a given phosphosite and category, the category score was defined as the maximum absolute standardised feature value among all features belonging to that category. This provides a per-site summary of the strongest functional signal within each class without assuming independence between individual features.

To assess whether phospho-dependent localisation was associated with a single dominant functional class or multiple contributing factors, we compared the highest and second-highest mean category scores for each phosphosite. The ratio between the strongest and second strongest category scores was used as a measure of dominance, where large ratios indicate a single dominant category and ratios close to one indicate comparable support from multiple categories. Category scores were visualised as heatmaps and dominant category assignments were summarised across sites to identify common functional classes associated with phosphorylation-dependent localisation.

### Analysis of subcellular transcriptomics and lncRNA relocalisation

Subcellular transcriptomics data from control and thapsigargin-treated U-2 OS cells were analysed using SANDLE. Each condition contained three biological replicates, each measured across eight subcellular fractions. Transcript annotations and marker labels were obtained from the previously processed LoRNA dataset. SANDLE was then applied to the six replicate-level datasets jointly, using marker annotations where available.

Differential localisation probabilities were computed using the paired experimental design, with three matched control and three matched thapsigargin-treated replicates. In parallel, a marker-free regression test was applied to the transcript fractionation profiles. Transcripts were retained only if all 48 measurements were present, corresponding to eight fractions across six replicate-level experiments. The marker-free test was then used to estimate the evidence for a condition-dependent change in fractionation profile, reported as an adjusted *q*-value and an effect size defined by the difference in residual sum of squares.

Marker transcripts were assigned a differential localisation probability of zero before downstream ranking to avoid interpreting training markers as candidate relocalisation events. Transcript annotations, including gene biotype and external gene name, were then joined to the SANDLE results. Differentially localised non-coding RNAs were defined using *q <*0.05. Results were visualised using volcano plots of localisation effect size against − log_10_(*q*), faceted by RNA biotype.

To identify lncRNAs showing a membrane-to-granule relocalisation pattern, we first selected lncRNAs passing the differential localisation threshold. For each candidate lncRNA, we compared the replicate-level abundance profiles between control and thapsigargin-treated cells. A transcript was classified as membrane-to-granule relocalising if it showed a consistent decrease in membrane-associated fraction 2 and a consistent increase in granule-associated fraction 6 after treatment across replicates, with mean control abundance in fraction 2 greater than 0.15. These transcripts were subsequently visualised as replicate-level fractionation profiles across all eight fractions.

For sequence-level analysis, Ensembl transcript identifiers were stripped of version suffixes and used to retrieve cDNA sequences from Ensembl BioMart. Sequence features were computed for each lncRNA, including transcript length, GC content, GC content in the first 100 nucleotides, total AUG count, AUG density, AUG count and density in the first 100 nucleotides, longest open reading frame length, and canonical G-quadruplex motif count. The longest open reading frame was defined as the longest sequence beginning with an AUG codon and terminating at an in-frame stop codon across all three reading frames. G-quadruplex motifs were counted using the canonical pattern (Wu *et al*., 2021):

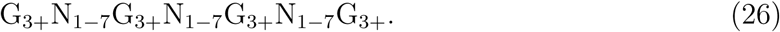

Sequence and profile-derived features were compared between membrane-to-granule lncRNAs and all other tested lncRNAs. Continuous features were compared using Wilcoxon rank-sum tests, followed by Benjamini–Hochberg correction across tested features. Enrichment of transcripts containing at least one G-quadruplex motif was assessed using Fisher’s exact test. Summary statistics were reported as group medians.

To assess which sequence features were most associated with membrane-to-granule relocalisation, we fitted logistic regression models with standardised predictors. The full model was

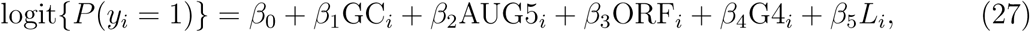

where *y_i_* indicates whether lncRNA *i* was classified as membrane-to-granule relocalising, GC*_i_* is transcript GC content, AUG5*_i_* is AUG density in the first 100 nucleotides, ORF*_i_* is the longest open reading frame length, G4*_i_* is the G-quadruplex motif count, and *L_i_* is transcript length. Single-predictor models containing either GC content or 5’ AUG density alone were also fitted and compared with the full model using Akaike’s information criterion.

### Trypanosome cross-species and life-cycle analysis

Spatial proteomics datasets from *Trypanosoma brucei* and *Trypanosoma congolense* were analysed to assess differential localisation across species and life-cycle stages. For cross-species comparisons, proteins were restricted to one-to-one orthologs identified using previously published orthology mappings (Emms and Kelly, 2019). This ensured that comparisons were performed on directly comparable protein sets.

SANDLE analysis was performed separately for within-species life-cycle comparisons and for cross-species comparisons. For life-cycle analyses within a species, marker annotations were used where available, and differential localisation probabilities were computed using the appropriate experimental design to account for biological replicates.

For cross-species comparisons, a marker-free analysis was used. This choice was motivated by the lack of reliable transferability of marker annotations across species, as marker proteins may not retain consistent localisation behaviour between *T. brucei* and *T. congolense*. To avoid introducing bias through potentially incorrect marker assignments, differential localisation was therefore assessed using the marker-free regression framework alone.

Replicate structure was incorporated by grouping replicate-level fractionation profiles within each condition. For life-cycle comparisons, replicates corresponded to biological repeats within each life-cycle stage. For cross-species comparisons, replicates were treated analogously across species to define the two conditions.

Differentially localised proteins were identified using SANDLE significance thresholds of FDR *<* 0.0001, corresponding to − log_10_(FDR) *>* 4, reflecting the large number of proteins tested and the desire for high-confidence cross-species comparisons. Where reported, differential localisation probabilities were used to further characterise the strength and consistency of spatial changes.

### Software and reproducibility

All analyses were performed in R (version 4.4.1). The SANDLE implementation is available in the BANDLE Bioconductor package.

## Supporting information

Supplementary manuscript

## Notes

### Competing Interest Statement

The authors have declared no competing interest.

### Funding

- Medical Research Council: MR/Y010078/1

