## Supplementary manuscript for "Robust analysis of comparative subcellular omics with complex designs"

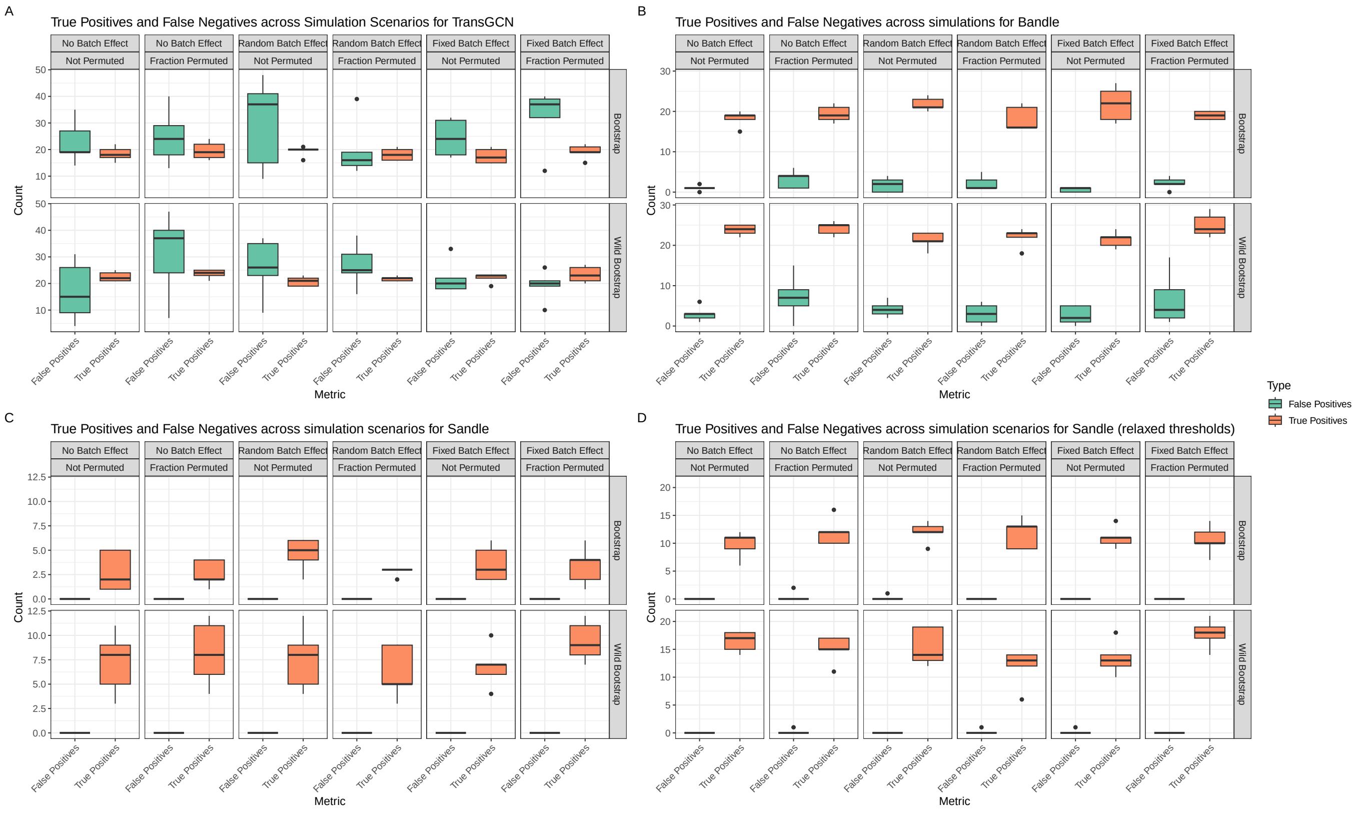


Figure S1: **Further analysis of simulation data** True positives and false positives across a range of scenarios for TransGCN, Bandle and Sandle

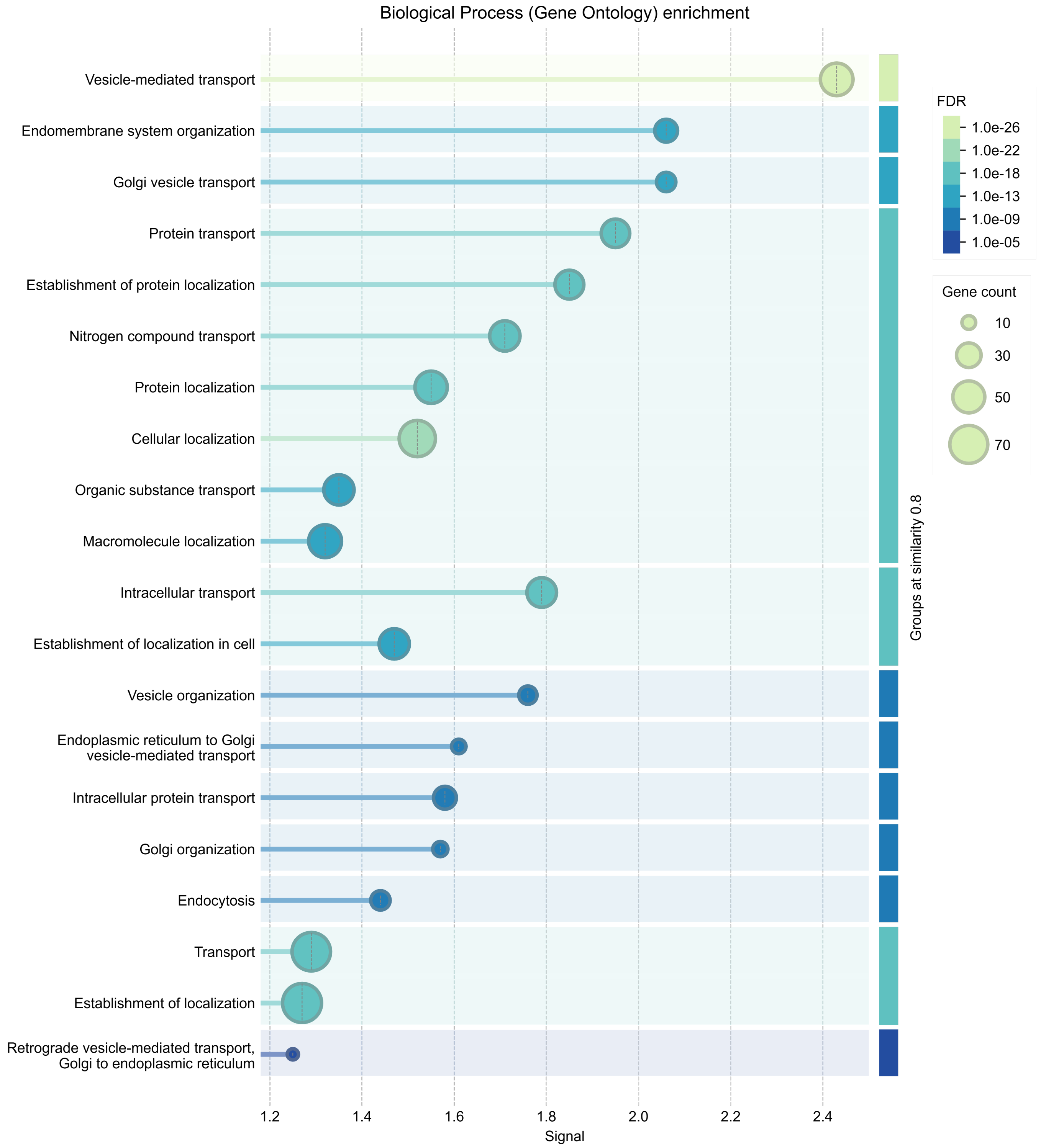


Figure S2: **GO enrichment of CHO versus MPC-11** Biological process enrichment on differentially localised proteins between CHO and MPC-11 cells.

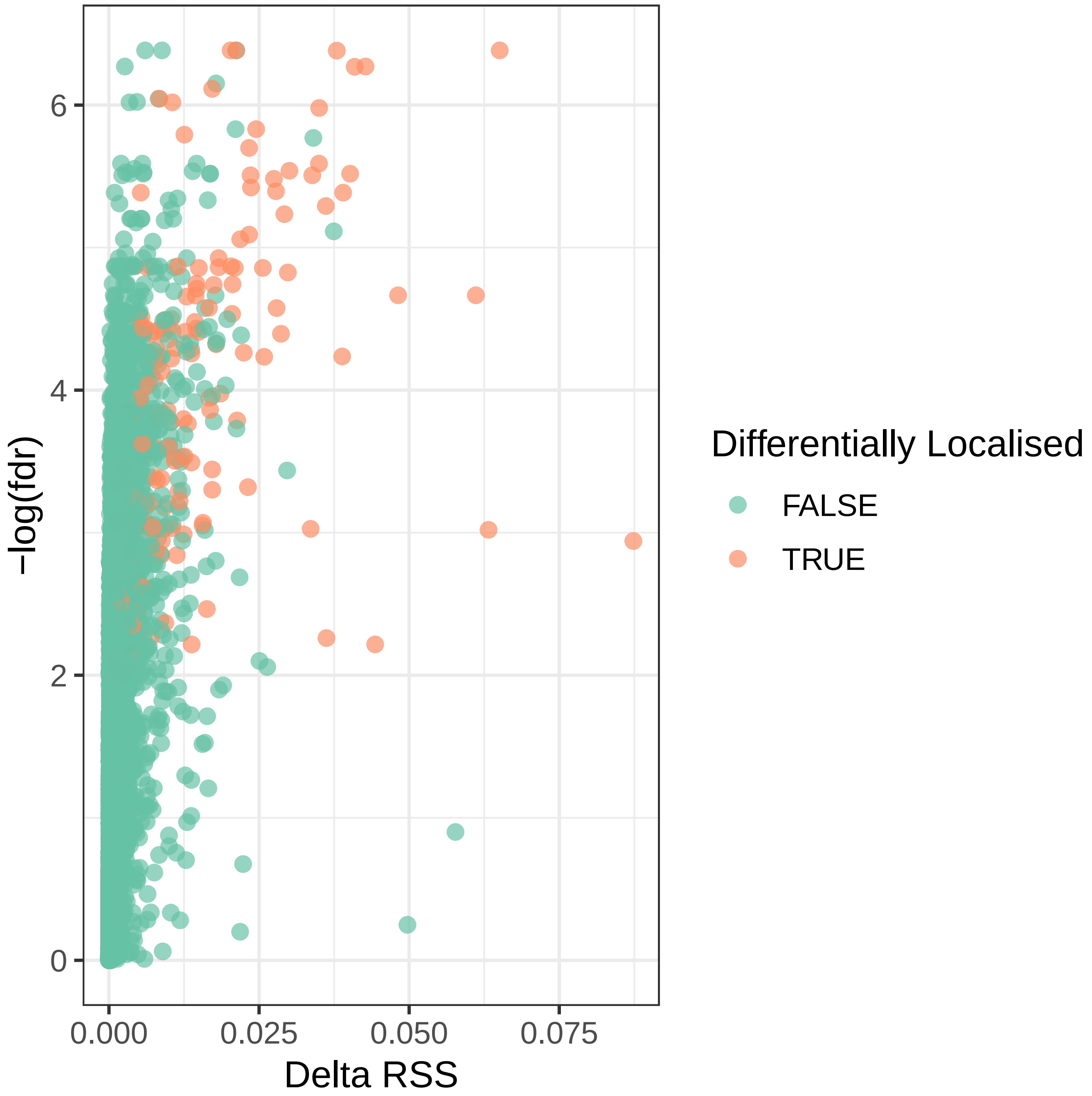


Figure S3: **Orre et al. Volcano Differential localisation** Volcano plots of proteins from the EGFR inhibition experiment. Highlighted proteins are detected at threshold FDR <0.01 and differential localisation probability 0.95.

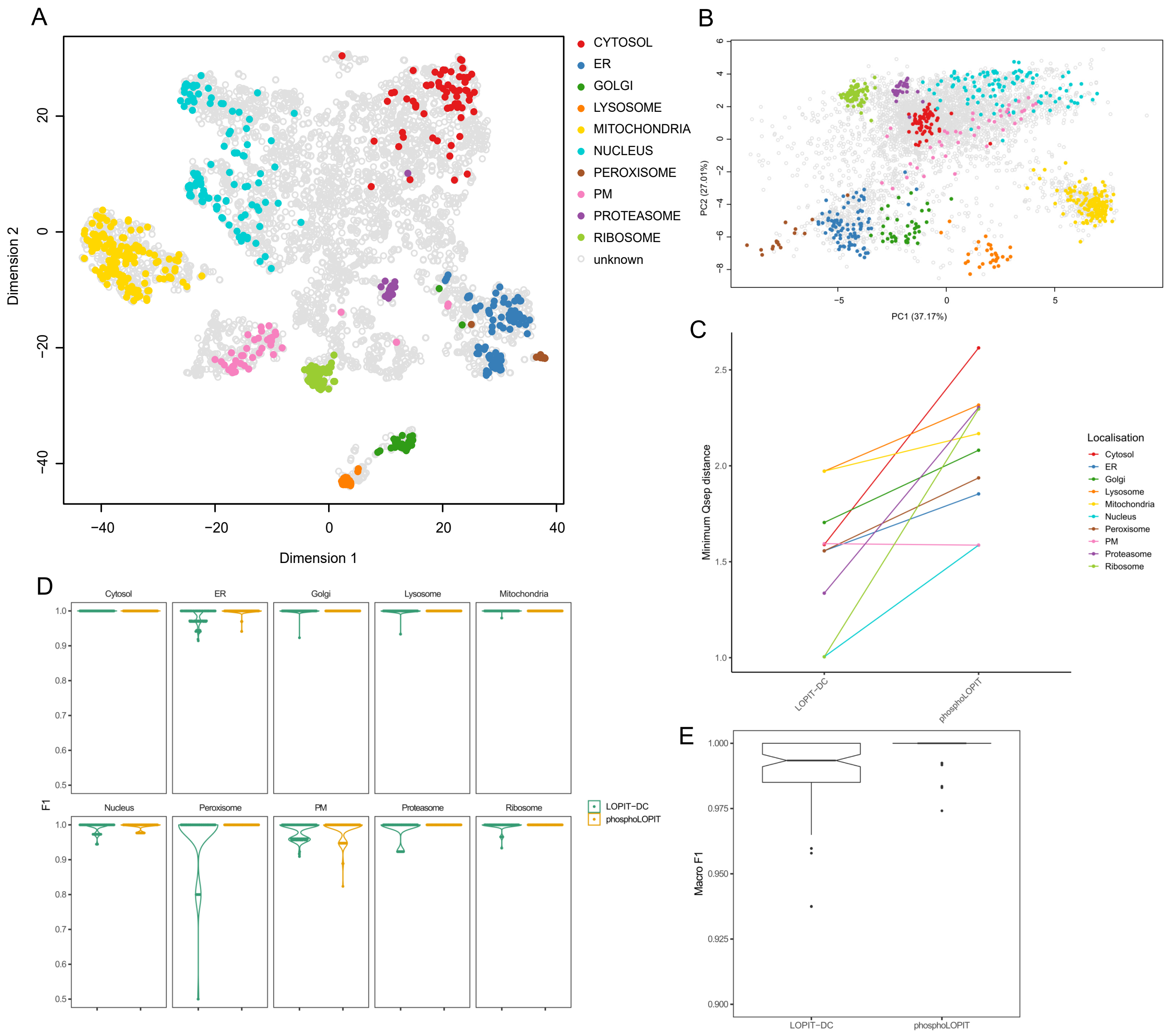


Figure S4: **Analysis and quality control of phosphoLOPIT** A) T-SNE plot of PhosphoLOPIT data show well separated subcellular compartments B) PCA plots of PhosphoLOPIT data showing well separated subcellular compartments C) QSEP scores showing generally improved resolution of PhosphoLOPIT compared with LOPIT-DC. D) Support vector machine F1 scores showing per organelle scores. Generally scores are higher for PhosphoLOPIT. E) Combined macro-F1 scores (all subcellular niches) showing improved classification accuracy for phosphoLOPIT proteins using a SVM

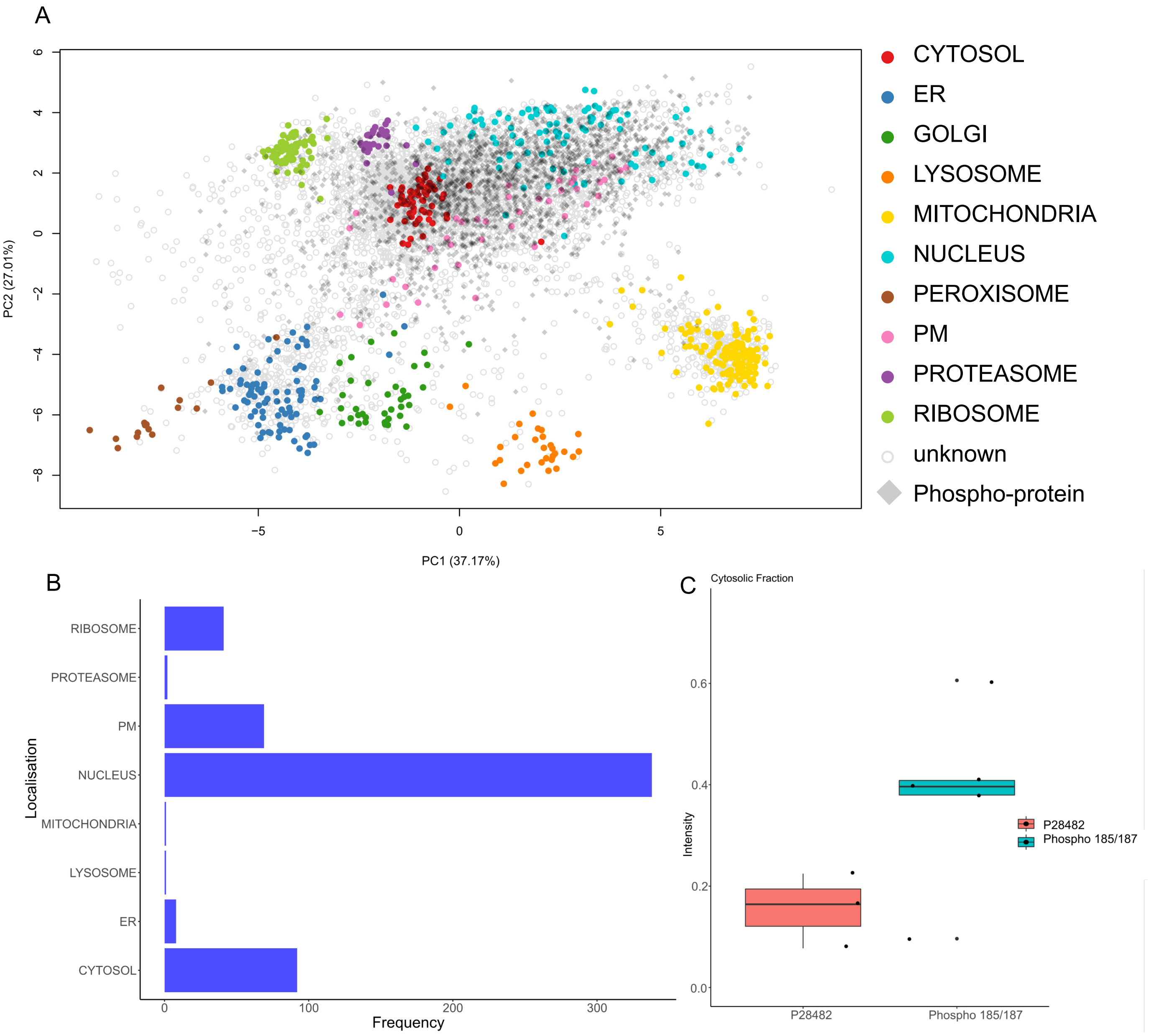


Figure S5: **Overview of phosphoLOPIT data** A) Phosphoproteins mapped onto total protein PCA plot. B) Overview of localisations of Phosphoproteins C) Cytosolic fraction intensities for ERK2

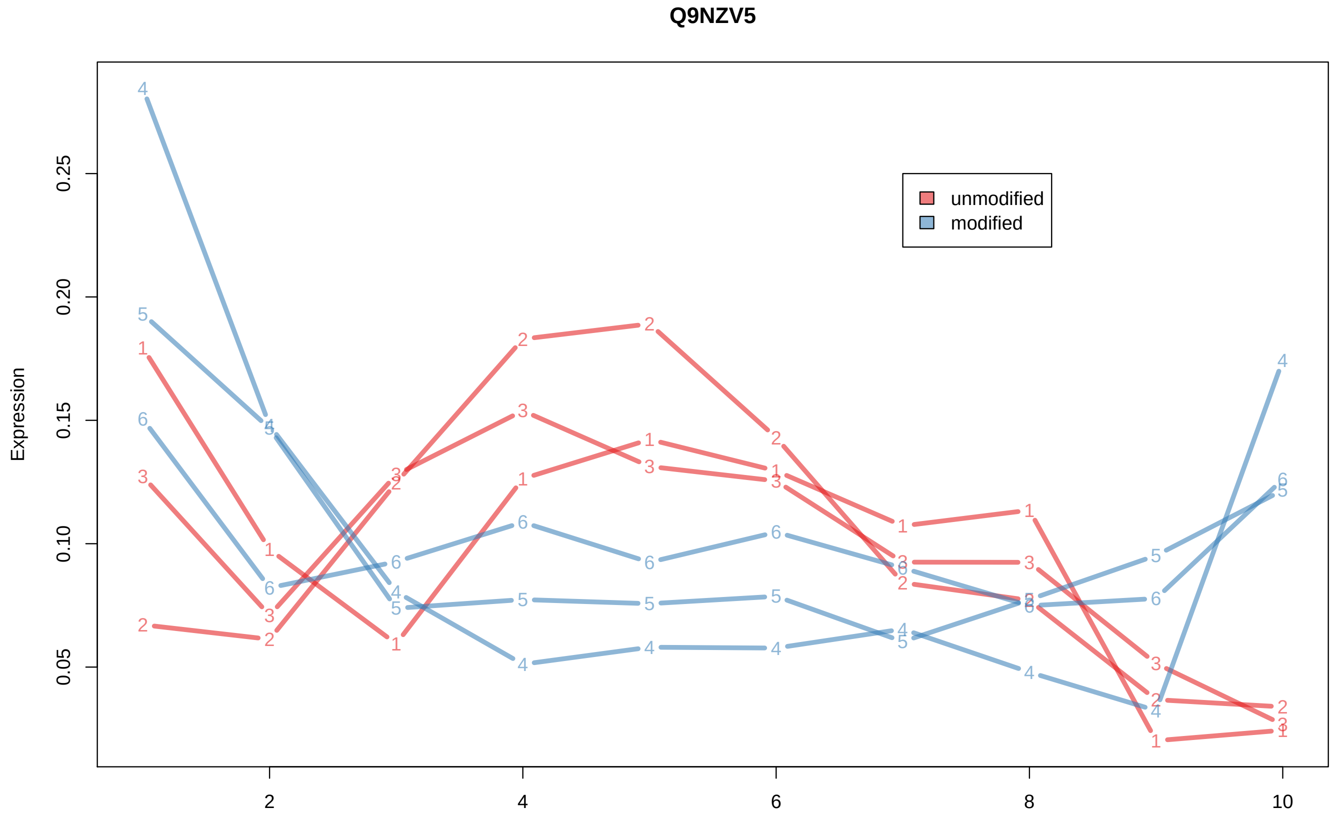


Figure S6: **Selenon abundance profiles** Abundance profiles of Selenon for modified and unmodified versions. Phosphorylated version is enriched in fraction 10 (cytosolic fraction).

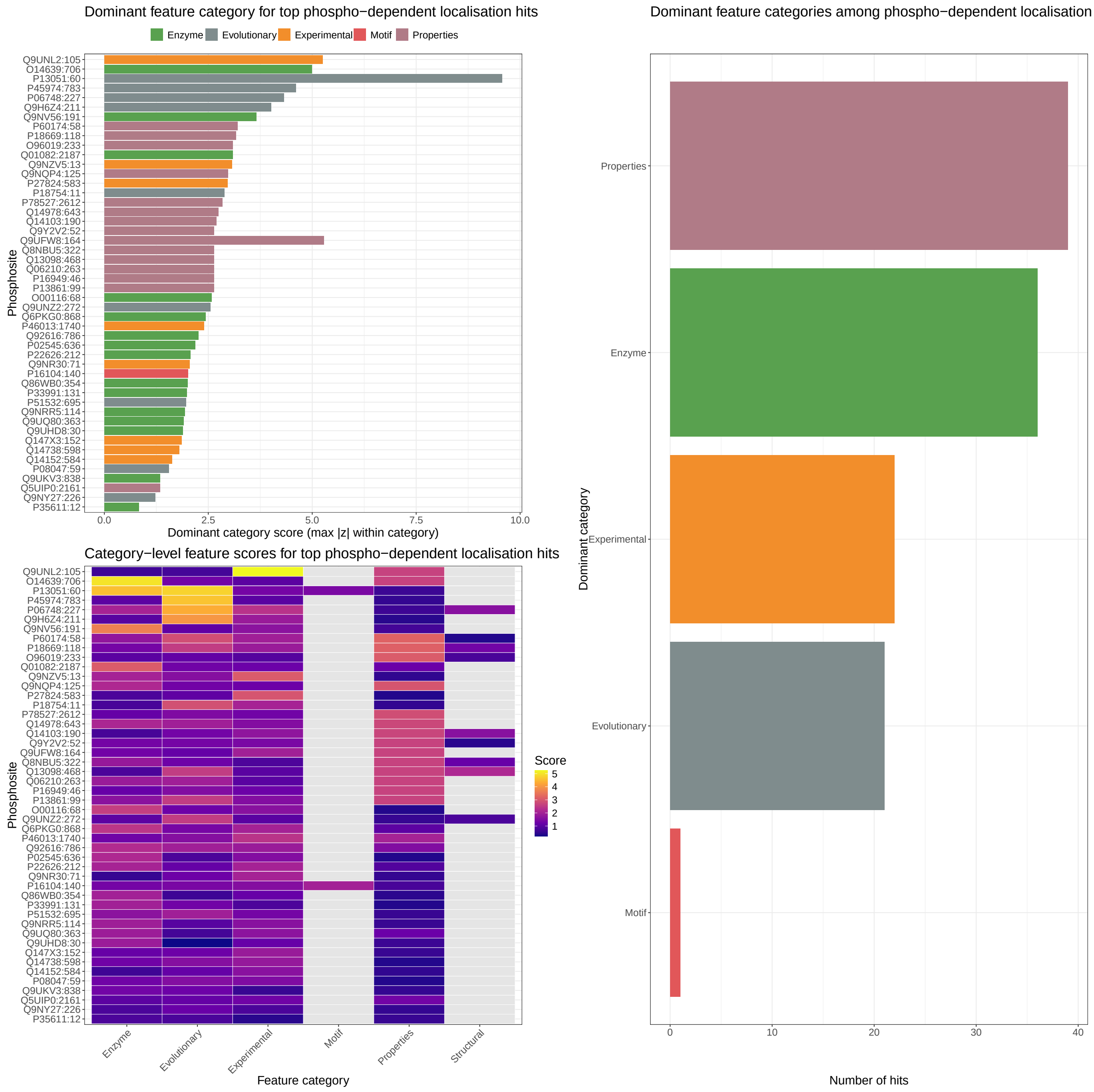


Figure S7: **Analysis of features predictive of Phospho-dependent localisation**. top-left) Dominant feature category from proteins with phospho-dependent localisations. bottom-left) Category-level feature scores for proteins with phosphorylation dependent localisation. Right) global overview of dominate feature categories

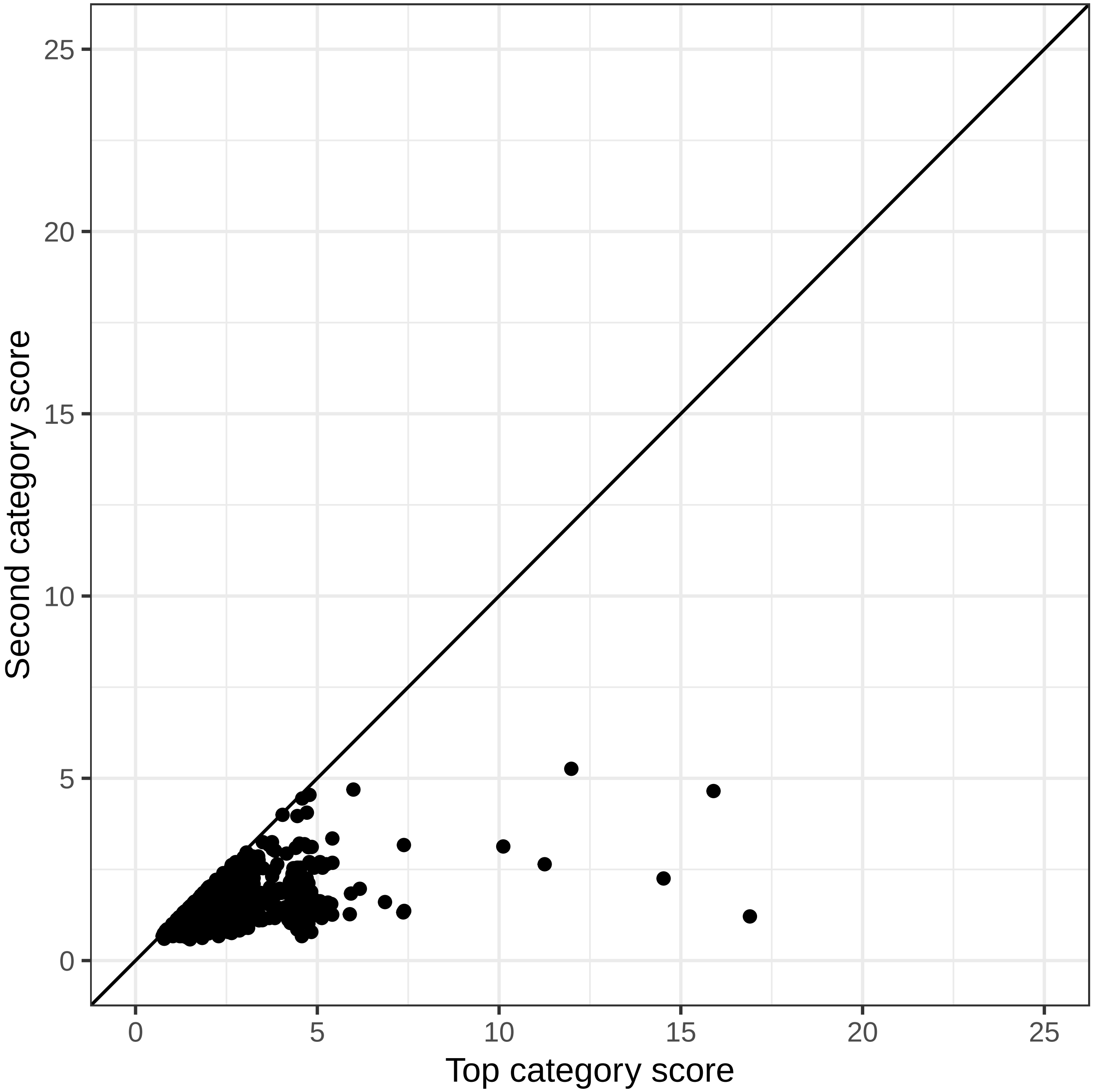


Figure S8: **Top versus second top score predictive of differential localisation**. Scatter plot of per protein highest score and second highest score for features that are predictive phospho dependent localisation

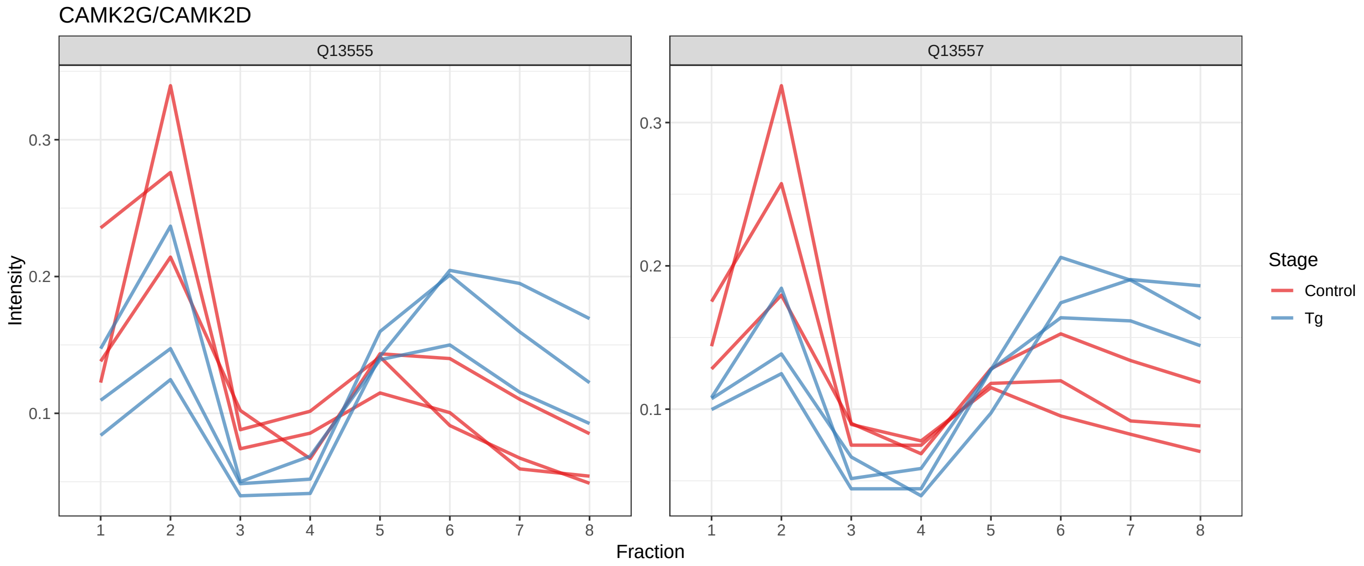


Figure S9: **CAMK2G and CAMK2D abundance profiles**

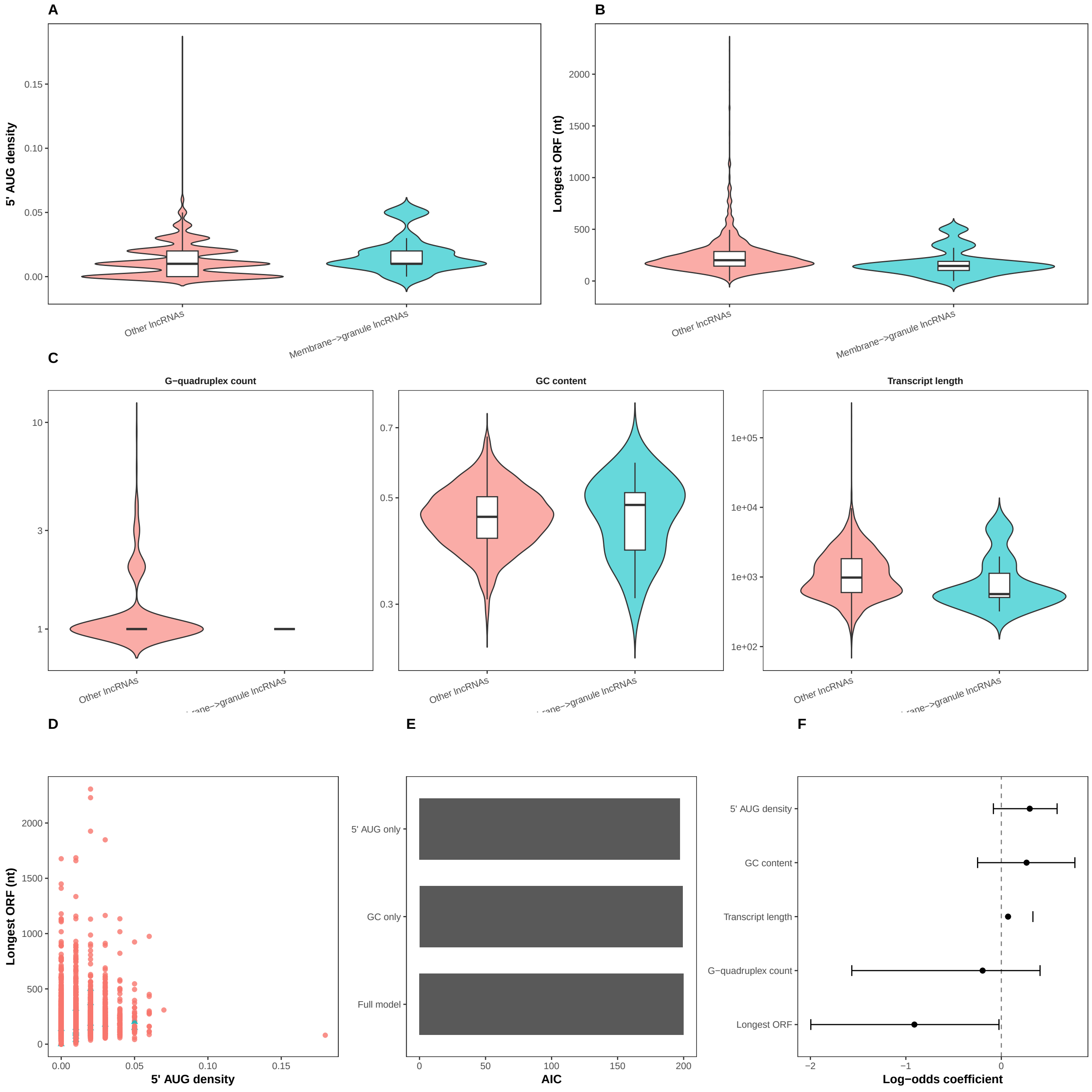


Figure S10: **Features of membrane to granule lncRNA re localisation** A) 5' AUG density B) longest ORF C) G-quandraplex count, GC content, Transcript length D) 5 AUG density versus longest ORF (uncorrelated) E) AIC for logistic models and full model combining all variables. F) log-odds coefficients for multivariate logistic models.
